# Cyclic AMP is a critical mediator of intrinsic drug resistance and fatty acid metabolism in *M. tuberculosis*

**DOI:** 10.1101/2022.07.07.499113

**Authors:** Andrew I. Wong, Tiago Beites, Kyle A. Planck, Shuqi Li, Nicholas C. Poulton, Kyu Rhee, Dirk Schnappinger, Sabine Ehrt, Jeremy Rock

**Affiliations:** Laboratory of Host-Pathogen Biology, The Rockefeller University, New York, New York, United States of America; Department of Microbiology and Immunology, Weill Cornell Medicine, New York, New York, United States of America; Division of Infectious Diseases, Department of Medicine, Weill Cornell Medicine, New York, New York, United States of America

## Abstract

Cyclic AMP (cAMP) is a ubiquitous second messenger that transduces signals from cellular receptors to downstream effectors. *Mycobacterium tuberculosis* (Mtb), the etiological agent of tuberculosis, devotes a considerable amount of coding capacity to produce, sense, and degrade cAMP. Despite this fact, our understanding of how cAMP regulates Mtb physiology remains limited. Here, we took a genetic approach to investigate the function of the sole essential adenylate cyclase in Mtb H37Rv, Rv3645. We found that lack of *rv3645* resulted in increased sensitivity to numerous antibiotics by a mechanism independent of substantial increases in envelope permeability. We made the unexpected observation that *rv3645* is conditionally essential for Mtb growth only in the presence of long-chain fatty acids, a host-relevant carbon source. A suppressor screen further identified mutations in the atypical cAMP phosphodiesterase *rv1339* that suppress both fatty-acid and drug sensitivity phenotypes in strains lacking *rv3645*. Using mass spectrometry, we found that Rv3645 is the dominant source of cAMP under standard laboratory growth conditions, that cAMP production is the essential function of Rv3645 in the presence of long-chain fatty acids, and that reduced levels of cAMP result in increased antibiotic susceptibility. Our work defines *rv3645* and cAMP as central mediators of intrinsic multidrug resistance and fatty acid metabolism in Mtb and highlights the potential utility of small molecule modulators of cAMP signaling.

## INTRODUCTION

*Mycobacterium tuberculosis* (Mtb) has evolved complex signaling and regulatory networks to sense and adapt to the diverse niches through which it transits during infection (Richard M Johnson and McDonough, 2018; Parish, 2014; Richard-Greenblatt and Av-Gay, 2017). The physiology that enables Mtb to adapt to and persist in the host can also decrease the effectiveness of antibiotics (Bellerose et al., 2020; Larrouy-Maumus et al., 2016). For example, hypoxia reduces Mtb respiratory capacity and activates the DosRST two-component system (Park et al., 2003). DosR induces a 48 gene regulon which ultimately slows growth, promoting survival under hypoxic conditions and tolerance to antibiotics that are more active against rapidly replicating bacteria (Galagan et al., 2013). In a second example, nutrient starvation results in the dephosphorylation of CwlM, a substrate of the eukaryotic-like protein serine/threonine kinase PknB (Boutte et al., 2016). Dephosphorylation reduces CwlM interaction with and activation of the peptidoglycan biosynthetic enzyme MurA, thereby reducing cell wall metabolism and promoting tolerance to starvation and antibiotics. Thus, the signaling and regulatory networks that facilitate Mtb survival under diverse physiologic conditions can also secondarily reduce the effectiveness of antibiotics. While several examples have been described, it is clear that numerous, poorly understood, adaptive mechanisms exist that promote Mtb survival while reducing the effectiveness of antibiotic therapy (Bellerose et al., 2020).

In addition to two-component systems and serine/threonine kinases, one of the most ubiquitous signal transduction modalities in Mtb are the adenylate cyclases. Adenylate cyclases sense environmental signals, either directly or indirectly, and transduce this signal into a cellular response by converting ATP into the small molecule second messenger 3’,5’-cyclic-AMP (cAMP) (Richard M. Johnson and McDonough, 2018). cAMP then binds to and alters the function of effector proteins like the transcription factor CRP (Stapleton et al., 2010), the protein lysine acetyltransferase Mt-Pat (Nambi et al., 2013), and numerous other potential cAMP-binding proteins (reviewed in Johnson and McDonough, 2018) to modify Mtb gene expression or gene product activity, thereby enabling Mtb to adapt to changes in its environment. Whereas the model bacteria *E. coli* encodes only one adenylate cyclase, the adenylate cyclase gene family has undergone dramatic expansion in mycobacteria. *M. avium, M. marinum*, and Mtb encode 12, 31, and at least 15 predicted adenylate cyclases (Shenoy and Visweswariah, 2006), respectively. The existence of so many adenylate cyclases in the Mtb genome presumably allows Mtb to integrate diverse environmental stimuli with downstream cellular responses by using cAMP as a second messenger. Mtb adenylate cyclases can be activated by a variety of host-relevant stimuli, including pH (Tews et al., 2005), bicarbonate (Cann et al., 2003), and fatty acids (Abdel Motaal et al., 2006), with the resulting increase in cAMP levels modulating both bacterial and host physiology (Agarwal et al., 2009). This expansion of adenylate cyclases is mirrored by an expansion of predicted cAMP phosphodiesterases, effectors, and binding proteins (**Supplemental Figure 1A**) – nearly 1% of the Mtb genome is predicted to produce, degrade, or interact with cAMP. However, despite the clear importance of cAMP signaling in mycobacteria, our knowledge of how adenylate cyclases and cAMP regulate Mtb physiology remain largely undefined.

To address this gap, we undertook a genetic study of the sole *in vitro* essential adenylate cyclase in Mtb H37Rv, *rv3645*. We found that lack of *rv3645* resulted in increased sensitivity to numerous antibiotics by a mechanism independent of substantial increases in envelope permeability. We further found that *rv3645* was conditionally essential for Mtb growth in the presence of long-chain fatty acids, a host-relevant carbon source, and identified mutations in the atypical cAMP phosphodiesterase *rv1339* that suppress this fatty-acid dependent essentiality. By leveraging gain-and loss-of-function alleles of *rv3645* and *rv1339* and small molecule-regulated cAMP production, our work defines Rv3645 and the ubiquitous second messenger cAMP as central mediators of intrinsic multidrug resistance and fatty acid metabolism in Mtb.

## RESULTS

### The essential adenylate cyclase Rv3645 contributes to intrinsic drug resistance in Mtb H37Rv

To identify genes and pathways that influence drug efficacy in Mtb, we previously screened a genome-wide CRISPRi library in Mtb strain H37Rv against a panel of diverse antibiotics (Li et al., 2022). This approach identified hundreds of Mtb genes whose inhibition altered bacterial fitness in the presence of partially inhibitory drug concentrations, including genes encoding the direct drug target and non-target hit genes. Amongst the non-target hit genes, we found that depletion of the predicted essential adenylate cyclase, *rv3645* (**Figure 1A**), sensitized Mtb to numerous antibiotics with unrelated mechanisms of action (**Figure 1B**). Interestingly, even though Mtb H37Rv encodes 15 putative adenylate cyclase genes, *rv3645* was the only adenylate cyclase gene whose knockdown resulted in increased drug sensitivity (**Supplemental Figure 1A**), demonstrating lack of substantial functional redundancy for adenylate cyclases under these culture conditions. Given the predicted essentiality of *rv3645* and the magnitude by which silencing of this gene sensitized Mtb to various antibiotics, we next sought to better characterize the role of this gene in Mtb physiology.

**Figure 1:**
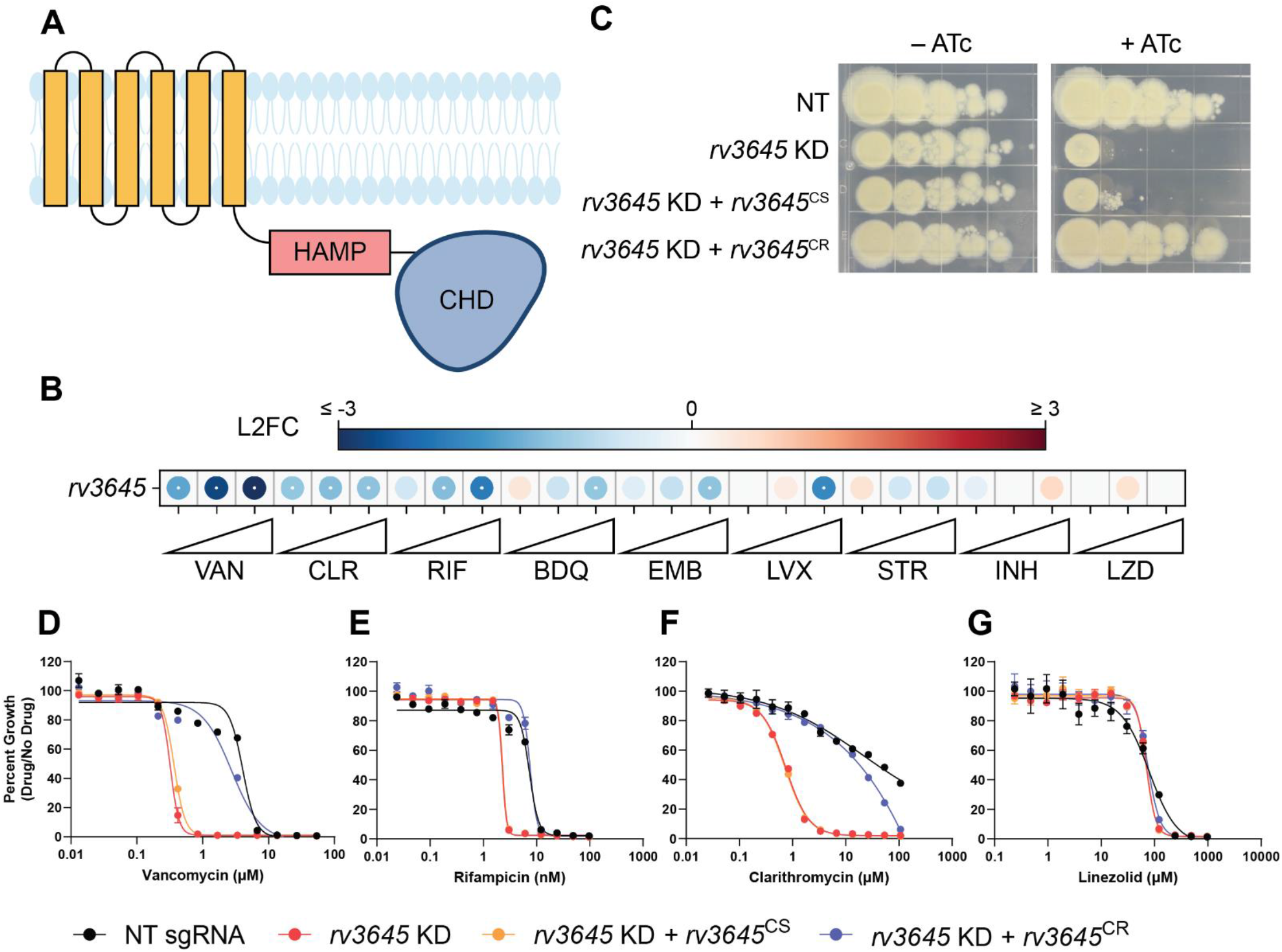
The adenylate cyclase *rv3645* is critical for intrinsic multidrug resistance in *M. tuberculosis*. (A) Predicted domain organization of Rv3645. N-terminal transmembrane helices, HAMP domain, and C-terminal cytosolic adenylate cyclase domain (CHD) are shown. HAMP = Histidine kinases, Adenylate cyclases, Methyl-accepting proteins and Phosphatases; CHD = Cyclase Homology Domain. (B) Feature-expression heatmap of *rv3645* from a 5-day CRISPRi library pre-depletion screen. The color of each circle represents the gene-level log2 fold change (L2FC); a white dot represents an FDR < 0.01 and a |L2FC| > 1. VAN = vancomycin; CLR = clarithromycin; RIF = rifampicin; BDQ = bedaquiline; EMB = ethambutol; LVX = levofloxacin; STR = streptomycin; INH = isoniazid; LZD = linezolid. (C) Growth of *rv3645* CRISPRi strains on 7H10-OADC agar. Columns represent 10-fold serial dilutions in cell number. NT = non-targeting sgRNA; KD = knockdown; CS = CRISPRi-sensitive; CR = CRISPRi-resistant. (D-G) Dose-response curves for (D) vancomycin, (E) rifampicin, (F) clarithromycin and (G) linezolid were measured against *rv3645* CRISPRi strains. Data represent mean ± SEM for technical triplicates and are representative of at least two independent experiments.

We first sought to validate the screen results. To do this, we cloned a single-guide RNA (sgRNA) targeting *rv3645* into an inducible CRISPRi plasmid that allows for targeted *rv3645* knockdown in the presence of anhydrotetracycline (ATc) (Rock et al., 2017). We also cloned complementation constructs expressing CRISPRi-sensitive or CRISPRi-resistant *rv3645* alleles (Wong and Rock, 2021). These *rv3645* complementation alleles differ only by silent mutations within the *rv3645* ORF that abrogate CRISPRi targeting in the resistant allele but do not affect the wild-type protein sequence. Consistent with prior screens (Bosch et al., 2021; DeJesus et al., 2017), knockdown of *rv3645* prevented growth on 7H10-OADC agar plates (**Figure 1C**). This growth defect was complemented by expressing a CRISPRi-resistant but not a CRISPRi-sensitive *rv3645* allele, demonstrating that growth inhibition was indeed a result of silencing *rv3645*. To validate the results of the CRISPRi chemical-genetic screen, we measured the minimum inhibitory concentrations (MICs) of the *rv3645* CRISPRi strains against a panel of antibiotics. Consistent with the CRISPRi screening results, *rv3645* knockdown sensitized Mtb to vancomycin, rifampicin, clarithromycin, bedaquiline, and meropenem but not other drugs (**Figure 1D-G; Supplemental Figure 1B-G**). These results validate that the sole essential adenylate cyclase Rv3645 contributes to intrinsic resistance of Mtb H37Rv to various antibiotics.

### Increased drug sensitivity in *rv3645* knockdown strains is not due to large increases in envelope permeability

The mycobacterial cell envelope serves as a permeability barrier that restricts access of antibiotics to their intracellular or periplasmic targets (Batt et al., 2020). The similarities between the chemical-genetic signatures of *rv3645* and envelope biosynthetic genes (**Figure 2A**) (Li et al., 2022) suggested that *rv3645* may contribute to intrinsic drug resistance by promoting envelope integrity in Mtb. To test this hypothesis, we used a fluorescent, BODIPY-conjugated analog of vancomycin (VANC-BODIPY) to monitor drug uptake. Vancomycin is a large, polar antibiotic for which disruption of the Mtb envelope is known to increase drug uptake and increase drug sensitivity. Surprisingly, despite the dramatic sensitization of *rv3645* knockdown strains to vancomycin (**Figure 1D**), *rv3645* knockdown strains showed only a modest increase in VANC-BODIPY uptake (**Figure 2B**) as compared to the positive control gene *mtrA*, encoding a two-component response regulator important for proper envelope biogenesis (Li et al., 2022). To ensure the discrepancy between *rv3645* MIC assays and drug uptake measurements was not due to the presence of the BODIPY conjugate (∼274 daltons), we next monitored drug uptake directly by mass spectrometry. Consistent with the VANC-BODIPY results, *rv3645* knockdown strains did not show elevated levels of vancomycin uptake (**Figure 2C**). Finally, we further monitored envelope permeability with an ethidium bromide uptake assay. Ethidium bromide is a hydrophobic, poorly permeable dye that fluoresces when it enters the cell and intercalates into Mtb genomic DNA. As with the vancomycin uptake assays, *rv3645* knockdown strains were not hyperpermeable to ethidium bromide (**Figure 2D**). Taken together, these results demonstrate that depletion of *rv3645* does not result in large increases in Mtb envelope permeability. Thus, Rv3645-mediated intrinsic drug resistance must occur primarily by some other mechanism.

**Figure 2:**
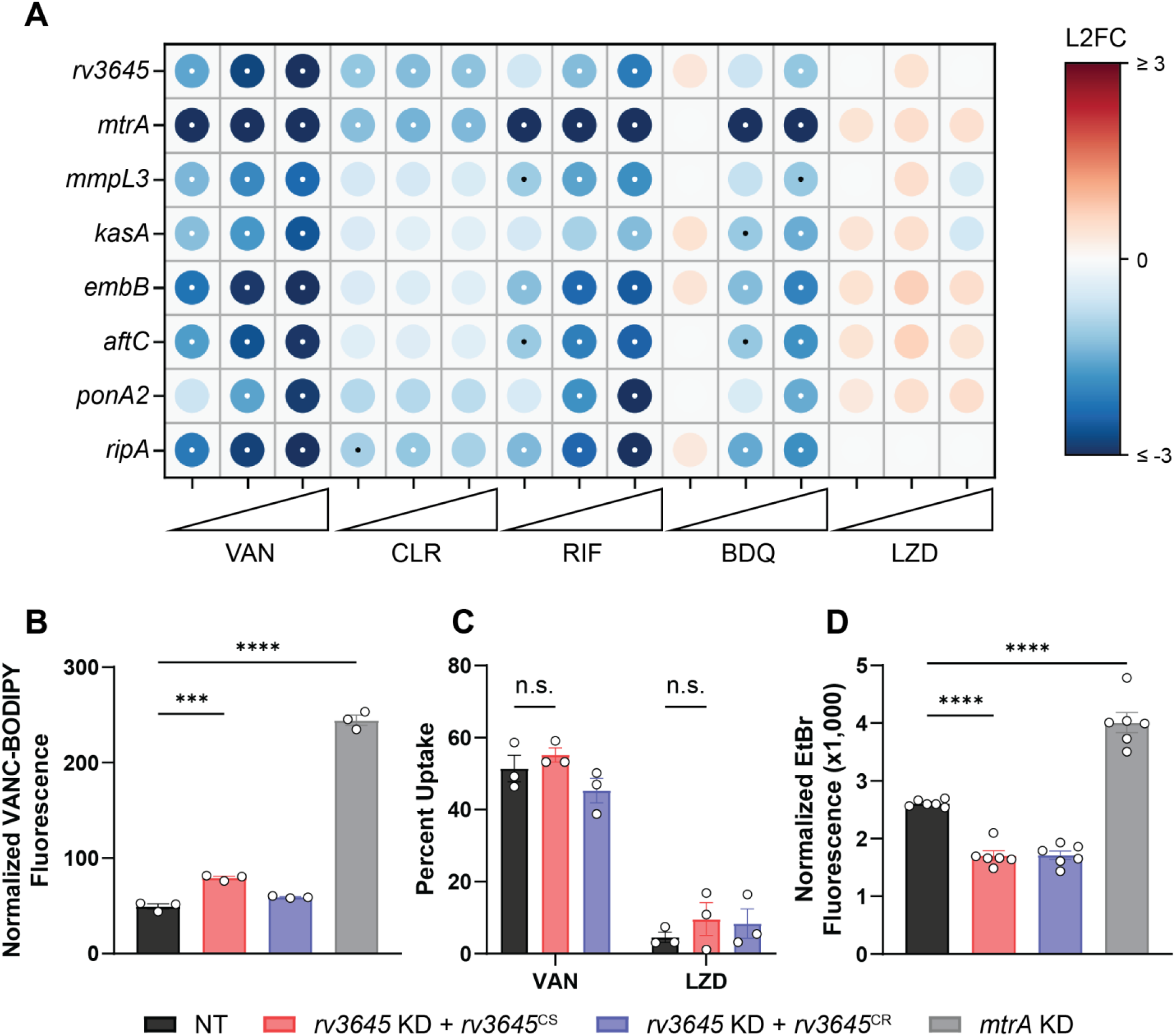
Increased drug sensitivity in *rv3645* knockdown strains is not due to large increases in envelope permeability. (A) Feature-expression heatmap of genes important for proper Mtb cell envelope biosynthesis from the 5-day CRISPRi library pre-depletion screen for select drugs. The color of each circle represents the gene-level L2FC; a white dot represents an FDR of < 0.01 and a |L2FC| > 1. VAN = vancomycin; CLR = clarithromycin; RIF = rifampicin; BDQ = bedaquiline; LZD = linezolid. (B) Vancomycin-BODIPY uptake of the indicated strains. Data represent mean ± SEM for three replicates. ***, p < 0.001; ****, p < 0.0001. Statistical significance was assessed by one-way ANOVA. (C) Quantification of vancomycin and linezolid uptake by mass spectrometry for the indicated strains. Data represent mean ± SEM for technical triplicates. n.s. = not significant. Statistical significance was assessed by two-way ANOVA (GraphPad Prism). (D) Ethidium bromide uptake of the indicated strains. NT = non-targeting sgRNA; KD = knockdown; CS = CRISPRi-sensitive; CR = CRISPRi-resistant. Data represent mean ± SEM for six replicates. ****, p < 0.0001. Statistical significance was assessed by one-way ANOVA.

### *rv3645* essentiality and contribution to intrinsic drug resistance is conditional on the presence of long-chain fatty acids

*rv3645* is essential for Mtb H37Rv growth in standard laboratory media (7H10 + OADC supplement: oleic acid, albumin, dextrose and catalase; **Figure 1C**). Curiously, when the same growth medium was instead supplemented without fatty acid (ADC), *rv3645* knockdown no longer inhibited growth (**Supplemental Figure 2A**), suggesting that *rv3645* is conditionally essential in the presence of oleic acid. Using fatty acid-free growth conditions, we were able to generate an *rv3645* deletion strain (Δ*rv3645*). Genetic identity was confirmed through whole genome sequencing. Plating of Δ*rv3645* in the presence or absence of oleic acid confirmed that *rv3645* is conditionally essential in the presence of this fatty acid (**Figure 3A**). To determine which other fatty acids may render *rv3645* essential, we measured MICs of fatty acids of increasing carbon chain lengths. Δ*rv3645* was uniquely sensitive to long chain fatty acids palmitic acid (C16:0), oleic acid (C18:1), and arachidonic acid (C20:4); but not short or medium chain fatty acids or cholesterol (**Figure 3B-D; Supplementary Figure 2B-G**). Notably, sensitivity was not observed towards the odd chain fatty acids propionic acid and valeric acid nor to cholesterol, ruling out propionate-derived toxicity as the source of the fatty acid sensitive growth phenotype (Eoh and Rhee, 2014). The fact that growth of Δ*rv3645* in the presence of long-chain fatty acids was not rescued by alternative carbon sources in the medium (glycerol and glucose) suggests that long-chain fatty acids are toxic to Δ*rv3645*, rather than Δ*rv3645* being unable to consume them. We next sought to determine if the contribution of *rv3645* to intrinsic drug resistance is also dependent on the presence of long chain fatty acids. As with the growth defect, drug sensitivity associated with Δ*rv3645* was also dependent on the presence of long chain fatty acids in the growth media (**Figure 3E-G)**. These results demonstrate that *rv3645* essentiality and contribution to intrinsic drug resistance are both conditional on the presence of long-chain fatty acids.

**Figure 3:**
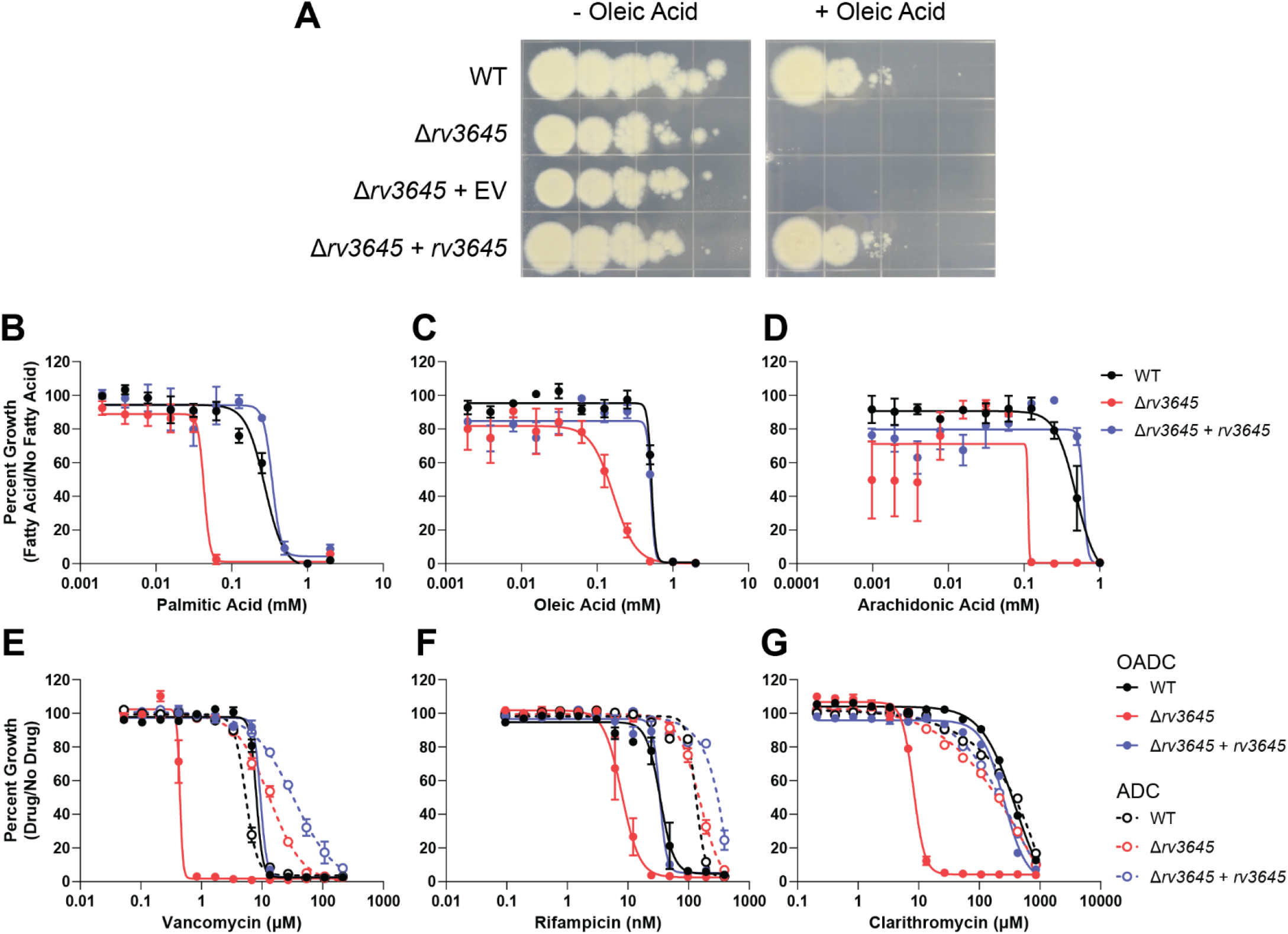
*rv3645* essentiality and contribution to intrinsic drug resistance is conditional on the presence of long-chain fatty acids. (A) Growth of Δ*rv3645* deletion strains on 7H10-ADC agar in the presence or absence of oleic acid. EV = empty vector. (B-D) Dose-response curves for fatty acids palmitic acid (B), oleic acid (C), and arachidonic acid (D). Data represent mean ± SEM for technical triplicates and are representative of at least two independent experiments. (E-G) Dose-response curves for vancomycin (E), rifampicin (F), clarithromycin (G) of the indicated strains grown in 7H9 with (OADC) or without (ADC) oleic acid.

### Loss-of-function of the atypical cAMP phosphodiesterase *rv1339* rescues fatty acid and drug sensitivity phenotypes of the Δ*rv3645* strain

Thus far, our results suggest that Rv3645 plays an important role in lipid metabolism in Mtb. To begin to interrogate how Rv3645 may contribute to lipid metabolism, we conducted a CRISPRi screen to identify suppressors of the fatty acid dependent growth defect of Δ*rv3645*. An ATc-inducible CRISPRi library consisting of 96,700 sgRNAs targeting 4,054/4,125 of all Mtb genes was transformed into Δ*rv3645* (**Figure 4A**) (Bosch et al., 2021). The resulting Δ*rv3645* CRISPRi library was then cultured on 7H10 agar in the presence of ATc and in the presence or absence of a toxic concentration of palmitic acid. As expected, growth was markedly reduced in the presence of palmitic acid. Colonies that grew in the presence of palmitic acid were expected to harbor sgRNAs that silence genes that contribute to fatty acid toxicity in Δ*rv3645*. Colonies were harvested from both culture conditions and their sgRNAs were PCR amplified, deep sequenced, and counted to compare sgRNA representation +/– palmitic acid. Hits genes were identified by MAGeCK (Li et al., 2014).

**Figure 4:**
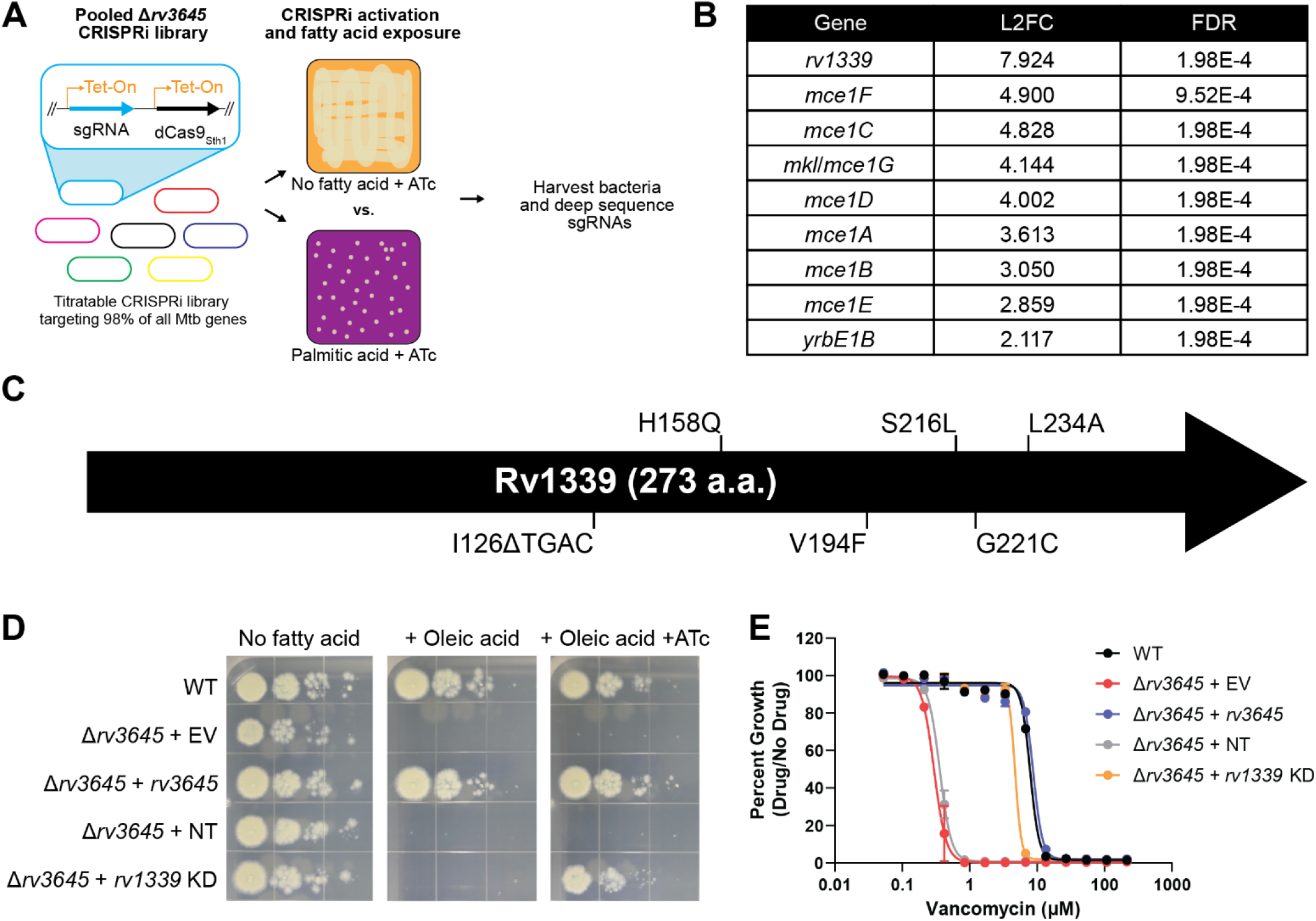
Loss-of-function of the atypical cAMP phosphodiesterase Rv1339 rescues fatty acid and drug sensitivity phenotypes of Δ*rv3645* strains. (A) Schematic of the Δ*rv3645* CRISPRi suppressor screen. First, an inducible genome-wide CRISPRi library was cloned into Δ*rv3645* Mtb. The CRISPRi library was then expanded before plating on 7H10-ADC agar supplemented with ATc in the presence or absence of an inhibitory concentration of palmitic acid (200 μM). Genomic DNA from surviving bacteria was prepared for sgRNA deep sequencing to identify genes whose inhibition permitted growth of an Δ*rv3645* strain in the presence of palmitic acid. (B) List of all enriched hit genes (log2 fold change (L2FC) > 2 and false discovery rate (FDR) < 0.01) from the suppressor screen described in panel (A). (C) Spontaneous suppressors of Δ*rv3645* oleic acid sensitivity were isolated and genomes sequenced. Identified mutations in *rv1339* are shown. (D) Growth of Δ*rv3645* CRISPRi suppressor strains. EV = empty vector; NT = non-targeting sgRNA; KD = knockdown. (E) Vancomycin dose-response curves of the indicated Δ*rv3645* strains. Data represent mean ± SEM for technical triplicates and are representative of at least two independent experiments.

The suppressor screen identified nine enriched genes, eight of which encode structural or catalytic subunits of the Mce1 transporter (**Figure 4B; Supplemental Data 1**). Mce1 has recently been shown to be an importer of fatty acids, including palmitic acid, in Mtb (Nazarova et al., 2019). Thus, Δ*rv3645* Mce1 knockdown strains likely fail to import palmitic acid, thereby allowing growth in the presence of this fatty acid. The fact that knockdown of the Mce1 transporter allows Δ*rv3645* to grow in the presence of palmitic acid suggests that the fatty acid toxicity phenotype of Δ*rv3645* is dependent on palmitic acid uptake and metabolism, rather than an uptake-independent toxicity mechanism such as fatty-acid dependent disruption of the Mtb envelope (Kengmo Tchoupa et al., 2022).

The top hit in the CRISPRi suppressor screen was the non-essential gene *rv1339* (**Figure 4B**). Consistent with loss of Rv1339 activity suppressing the fatty acid sensitive phenotype of an Δ*rv3645* strain, isolation of spontaneous suppressors of Δ*rv3645* grown in the presence of oleic acid identified five unique mutations in *rv1339*, including one frameshift mutation (**Figure 4C**). We confirmed that CRISPRi knockdown of *rv1339* resulted in rescue of Δ*rv3645* fatty acid sensitivity with individual strains (**Figure 4D**). To test whether loss of Rv1339 activity also suppresses the fatty acid dependent drug sensitivity phenotype, we performed MIC assays in a Δ*rv3645 rv1339* knockdown strain. Indeed, *rv1339* knockdown rescued the vancomycin sensitivity phenotype of the Δ*rv3645* strain (**Figure 4E**).

Intriguingly, Rv1339 was recently reported to be an atypical cAMP phosphodiesterase (Thomson et al., 2022). While Rv3645 is the sole essential adenylate cyclase in H37Rv, it is possible that one or more of the other 14 additional adenylate cyclase homologues encoded in the genome could also be synthesizing cAMP under these growth conditions. In this case, knockdown of the cAMP-degrading enzyme Rv1339 could restore cAMP levels in an Δ*rv3645* strain. These results strongly implicate cAMP levels in coordinately regulating fatty acid metabolism and intrinsic drug resistance in Mtb.

### The second messenger cAMP is a critical mediator of fatty acid metabolism and multidrug intrinsic resistance in Mtb

To test the hypothesis that cAMP coordinately regulates fatty acid metabolism and intrinsic drug resistance in Mtb, we first sought to test whether *rv3645* mutants incapable of synthesizing cAMP could complement these phenotypes. We cloned a *rv3645* allele with a point mutation in a metal-coordinating residue known to be essential for adenylate cyclase catalytic activity (Linder et al., 2002). The Rv3645 adenylate cyclase domain point mutant was unable to complement oleic acid and vancomycin sensitivity phenotypes (**Figure 5A, B**). We confirmed by western blot that the catalytic mutant was expressed at wild-type levels, and thus we attribute the lack of complementation specifically to the loss of cAMP synthesis specifically (**Supplementary Figure 3**).

**Figure 5:**
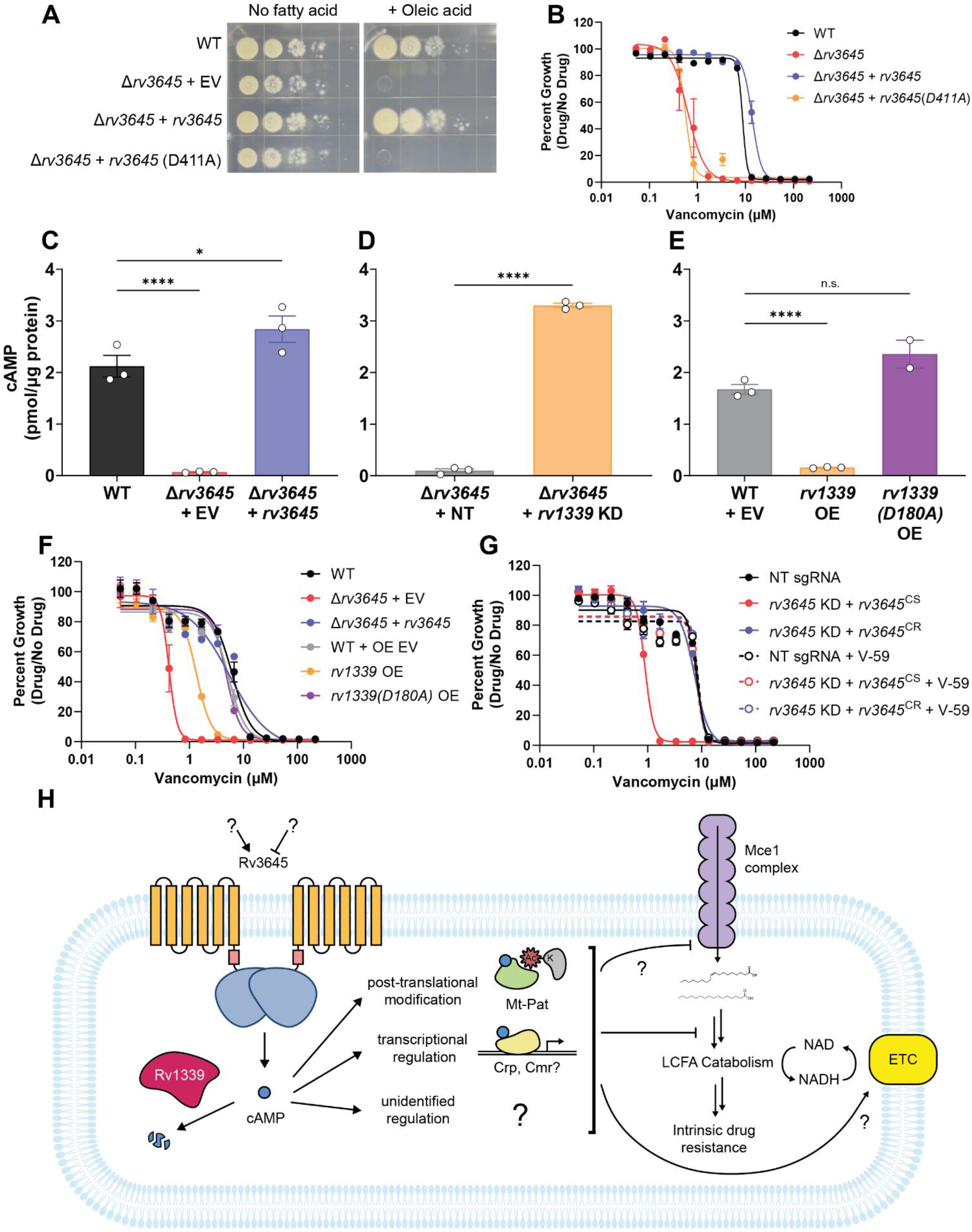
The second messenger cAMP is a critical mediator of fatty acid metabolism and multidrug intrinsic resistance in *M. tuberculosis*. (A) Growth on 7H10-ADC +/-oleic acid of indicated Δ*rv3645* strains expressing a *rv3645* adenylate cyclase catalytic mutant (D411A) (B) Vancomycin dose-response curves of indicated CRISPRi strains with adenylate cyclase *rv3645* (D411A) catalytic mutant defective in cAMP production. Data represent mean ± SEM for technical triplicates. (C-E) cAMP measurement of the indicated strains. D180A is a catalytically dead *rv1339* allele. EV = empty vector; OE = over-expressed. Data represent mean +/-SEM for technical triplicates. n.s. = not significant; *, p < 0.05; ****, p < 0.0001. Statistical significance was assessed by one-way ANOVA (GraphPad Prism). (F) Vancomycin dose-response curves of *rv3645* deletion mutants overexpressing *rv1339*. OE EV = overexpression empty vector. Data represent mean ± SEM for technical triplicates and are representative of at least two independent experiments. (G) Vancomycin dose-response curves of the *rv3645* CRISPRi mutants grown in the presence (dotted lines) or absence (solid lines) of the adenylate cyclase Rv1625 agonist V-59. NT = non-targeting sgRNA; KD = knockdown; CS = CRISPRi-sensitive; CR = CRISPRi-resistant. (H) Model for involvement of cAMP in lipid metabolism and intrinsic drug resistance. Under standard lab culture conditions (7H9/7H10 media), Rv3645 produces the majority of cAMP in Mtb. cAMP regulates physiological processes through binding to transcription factors, post-translational modification enzymes, and potentially other unknown mediators. Regulation through one or more of these mechanisms affects the fate of mce1-imported long-chain fatty acids (LCFA), with downstream effects on intrinsic drug resistance.

To validate that loss of Rv3645 reduces intracellular cAMP levels in Mtb grown in 7H9-OADC, we quantified cAMP levels by mass spectrometry. A 31-fold reduction in cAMP was observed in the *rv3645* deletion strain (**Figure 5C**) that could be rescued by knockdown of *rv1339* (**Figure 5D**). Conversely, overexpression of *rv1339* in an otherwise wild-type background led to an 11-fold reduction in cAMP levels (**Figure 5E**). Overexpression of a catalytically dead *rv1339* allele did not reduce cAMP levels (**Figure 5E**) (Thomson et al., 2022). Consistent with the role of cAMP in modulating drug sensitivity, overexpression of *rv1339* but not the catalytically dead *rv1339* allele resulted in increased vancomycin sensitivity in Mtb (**Figure 5F**). To demonstrate that modulating cAMP levels is sufficient to regulate Mtb H37Rv antibiotic sensitivity, we treated *rv3645* knockdown strains with V-59, a small molecule agonist of the adenylate cyclase Rv1625c that results in constitutive cAMP production (Wilburn et al., 2022). Addition of V-59 rescued the vancomycin sensitivity of *rv3645* knockdown strains (**Figure 5G**). Taken together, these data are consistent with a critical role for cAMP in regulating fatty acid metabolism and intrinsic drug resistance in Mtb.

## DISCUSSION

Our work defines *rv3645* and cAMP as central mediators of intrinsic multidrug resistance and fatty acid metabolism in Mtb H37Rv. *rv3645* knockdown resulted in increased sensitivity to antibiotics with diverse targets by a mechanism largely independent of increases in cell envelope permeability. Surprisingly, we found that *rv3645* was only essential in H37Rv in the presence of long-chain fatty acids. Suppression of long-chain fatty acid sensitivity was conferred by loss of the Mce1 transporter, presumably reflecting reduced fatty acid uptake, or inactivation of the atypical cAMP phosphodiesterase Rv1339. Consistent with these genetic results, using mass spectrometry we found that Rv3645 is the dominant source of cAMP under standard laboratory conditions in H37Rv, despite this strain encoding 14 additional adenylate cyclases in its genome. We propose naming Rv3645 “MacE” for major adenylate cyclase enzyme. Using gain-and loss-of-function alleles and small molecule-regulated cAMP production, we show that reduced cAMP levels are associated with fatty acid and drug sensitivity.

Why does lack of *rv3645* and lowered cAMP levels make Mtb more sensitive to fatty acids and drugs? Previous work established links between cAMP and fatty acid metabolism. Mt-Pat is a lysine acetyltransferase that is activated by cAMP (Nambi et al., 2013). Activated Mt-Pat acetylates/propionylates multiple enzymes involved in fatty acid catabolism, including ten FadD paralogs and acetyl-CoA synthetase, thereby inhibiting AMP-ligase activity and oxidation of fatty acids to acetyl-CoA. Inactivation of Mt-Pat, like *rv3645*, makes Mtb more sensitive to fatty acids (Nambi et al., 2013; Rittershaus et al., 2018), although the fatty acid sensitivity phenotype is more severe for loss of *rv3645* as evidenced by the non-essentiality of Mt-Pat under standard laboratory culture conditions (DeJesus et al., 2017). This work suggests that reduced cAMP levels, as in the absence of *rv3645*, may lead to overactive fatty acid catabolism.

In a series of related findings, Sassetti and colleagues screened Mtb transposon mutant libraries and identified Mt-Pat mutants as strongly attenuated under hypoxia (Rittershaus et al., 2018). In the absence of Mt-Pat, it was hypothesized that Mtb fails to downregulate fatty acid catabolism under hypoxic conditions, leading to continual flux of acetyl-CoA through oxidative TCA metabolism. Consistent with this interpretation, the same screen identified transposon mutants in the Mce1 transporter promoted fitness in hypoxia. Under hypoxia, the TCA cycle is thought to preferentially work in the reductive direction to regenerate NAD and prevent the accumulation of NADH in the absence of a terminal electron acceptor (Eoh and Rhee, 2013; Watanabe et al., 2011). As both the oxidative branch of the TCA cycle and fatty acid β-oxidation produce NADH, Δ*mt-pat* under hypoxic conditions becomes redox imbalanced and depletes NAD. Also potentially consistent with this interpretation, re-analysis of the hypoxia TnSeq data identifies *rv1339* as the top resistance promoting hit (Rittershaus et al., 2018). Elevated cAMP levels in the absence of *rv1339* could augment Mt-Pat activity and reduce fatty acid catabolism, potentially reduce fatty acid uptake (Nazarova et al., 2019), and drive expression of malate dehydrogenase (Gazdik and McDonough, 2005), an enzyme critical for the reductive TCA cycle under hypoxia (Rittershaus et al., 2018).

Finally, recent observations further suggest a link between cAMP and the electron transport chain. Isolation of spontaneous resistant mutants to a series of novel inhibitors of the QcrB subunit of the cytochrome *bc1-aa3* oxidase repeatedly identify mutations in *rv1339* (Chandrasekera et al., 2017; O’Malley et al., 2018; Shelton et al., 2021). How loss of *rv1339* and elevated cAMP promotes resistance to QcrB inhibitors remains to be defined, but in principle could be achieved by elevated cAMP levels increasing electron transport chain activity, for example by increasing expression of the alternative terminal oxidase cytochrome *bd* (Ko and Oh, 2020).

Collectively, ours and published results lead to the following working model (**Figure 5H**). Loss of *rv3645* reduces cAMP levels which reduces the activity of multiple cAMP effectors, including Mt-Pat. That no single annotated or predicted cAMP effector individually recapitulates the phenotypes observed with *rv3645* indicates that the physiological effects of reduced cAMP levels in Δ*rv3645* are the consequence of altered activity of multiple effectors (**Supplemental Figure 1**). Potential downstream cAMP effector proteins include transcription factors like CRP and Cmr, transporters, phospholipases, and others (Richard M. Johnson and McDonough, 2018), highlighting the potentially pleiotropic consequences of modulating cAMP levels. Reduced activity of cAMP-responsive proteins, including Mt-Pat, may increase fatty acid uptake and catabolism (Nazarova et al., 2019) and oxidative TCA metabolism while simultaneously reducing electron transport chain activity (Ko and Oh, 2020). This derangement may ultimately lead to redox imbalance, reduced growth, and increased sensitivity to antibiotics through yet to be defined mechanisms. Much work remains to be done to test the predictions generated by this model. Moreover, it is known that host-associated signals like low pH can activate Mtb adenylate cyclases and that macrophage infection results in a burst of bacterial cAMP production (Bai et al., 2009; Tews et al., 2005), and thus it is likely that some of the up to 15 additional adenylate cyclases encoded in the Mtb genome are active in the diverse niches encountered by Mtb during infection. Whether cAMP produced under these diverse host-associated conditions also coordinates fatty acid metabolism and intrinsic multidrug resistance remain to be investigated.

We show that the sole *in vitro* essential adenylate cyclase in H37Rv, Rv3645, links fatty acid metabolism and intrinsic multidrug resistance through the production of cAMP. Rv3645 is predicted to be composed of six transmembrane helices and cytosolic HAMP (Histidine kinases, Adenylate cyclases, Methyl-accepting proteins and Phosphatases) and cyclase homology domains. The presence of a HAMP domain, which typically transmits conformational changes from a periplasmic or transmembrane ligand binding domain to a cytoplasmic signaling domain (Hulko et al., 2006), suggests that Rv3645 senses an extracellular signal and transduces this signal into cytoplasmic cAMP production. The nature of this signal remains to be determined. The adenylate cyclase Rv1625c has recently been shown to be important for cholesterol catabolism, although interestingly this phenotype was independent of the Rv1625c cyclase homology domain (Wilburn et al., 2022). In a screen for compounds that disrupt cholesterol catabolism, VanderVen and colleagues discovered V-59, an Rv1625c agonist analogous to the eukaryotic adenylate cyclase activator forskolin (VanderVen et al., 2015; Wilburn et al., 2022). V-59 results in constitutive activation of Rv1625c and cAMP production and, for reasons yet unknown, blocks cholesterol catabolism. Thus, it would appear proper metabolic function of Mtb grown on fatty acids and cholesterol requires “just the right amount” of cAMP: too little and Mtb cannot grow in the presence of fatty acids, too much and Mtb cannot catabolize cholesterol. Moreover, our results and those investigating combination drug treatment with V-59 (Wilburn et al., 2022) suggest that deficient or elevated cAMP production may potentiate combination drug therapy.

Interestingly, whereas cAMP promotes intrinsic multidrug resistance in Mtb, this relationship may not be conserved or in some cases may even be reversed in other bacteria. Large-scale chemical genomic screening of cAMP-deficient Δ*cya E. coli* did not identify any significant differences in antibiotic susceptibility (Nichols et al., 2011), although earlier studies found that Δ*cya* in *E. coli* and *S. typhimurium* promotes fosfomycin resistance by reducing expression of the GlpT and UhpT uptake systems (Shiver et al., 2016; Silver, 2017). In uropathogenic *E. coli*, cAMP was also found to be a negative regulator of persistence (Molina-Quiroz et al., 2018). Δ*cya* upregulated the oxidative stress response and SOS-dependent DNA damage repair and promoted survival to beta-lactam antibiotics. Thus, it will be interesting to examine the relationship between cAMP and drug efficacy across diverse bacterial species.

cAMP is a ubiquitous but poorly understood second messenger in Mtb. Mtb devotes a considerable amount of coding capacity to produce, sense, and degrade cAMP. Here, we reveal adenylate cyclase Rv3645 as the dominant source of cAMP under standard laboratory growth conditions. cAMP levels are coordinately regulated by Rv3645 and the atypical phosphodiesterase Rv1339. cAMP produced by Rv3645 is critical for Mtb growth on long-chain fatty acids, a host-relevant carbon source, and for intrinsic multidrug resistance, highlighting the potential utility of small molecule modulators of this second messenger to control Mtb infection (Wilburn et al., 2022).

## MATERIALS AND METHODS

### BACTERIAL STRAINS

Mtb strains are derivatives of H37Rv. *E. coli* strains are derivatives of DH5alpha (NEB). *M. smegmatis* strains are derivatives of mc^2^155 *groEL1*Δ*C* (Noens et al., 2011)

### MYCOBACTERIAL CULTURES

Mtb was grown at 37°C in Difco Middlebrook 7H9 broth or on 7H10 agar supplemented with 0.2% glycerol (7H9) or 0.5% glycerol (7H10), 0.05% Tween-80, 1X oleic acid-albumin-dextrose-catalase (OADC) and the appropriate antibiotics, unless otherwise specified. Media for the Δ*rv3645* strain and strains to be tested for fatty acid sensitivity or fatty acid-dependent phenotypes were similarly prepared except 0.05% tyloxapol was used instead of Tween-80, and fatty acid free albumin-dextrose-catalase (ADC) was used instead of OADC. Where required, antibiotics or small molecules were used at the following concentrations: kanamycin at 20 μg/mL; anhydrotetracycline (ATc) at 100 ng/mL, hygromycin at 50 μg/mL, and zeocin at 20 μg/mL. Mtb cultures were grown standing in tissue culture flasks (unless otherwise indicated) at 37°C, 5% CO_2_. Fatty acid sensitivity testing on 7H10 agar was conducted with 500 μM oleic acid.

*M. smegmatis* was grown at 37°C in similarly supplemented 7H9 broth or 7H10 agar except ADC was used instead of OADC.

### DOMAIN PREDICTION

Rv3645 domain prediction was conducted using the Conserved Domain Database (https://www.ncbi.nlm.nih.gov/Structure/cdd/wrpsb.cgi). Transmembrane helices were predicted using TMHMM (https://services.healthtech.dtu.dk/service.php?TMHMM-2.0).

### ANTIBACTERIAL ACTIVITY AND FATTY ACID SENSITIVITY MEASUREMENTS

All antibiotics were dissolved in DMSO (VWR V0231) and dispensed using an HP D300e Digital Dispenser in a 384-well plate format. DMSO did not exceed 1% of the final culture volume and was maintained at the same concentration across all samples. CRISPRi strains were growth-synchronized and pre-depleted in the presence of ATc (100 ng/mL) for 4 days prior to assay for MIC analysis. Cultures were then back diluted to a starting OD_600_ of 0.05 in 7H9-OADC, and 50 μL of cell suspension was plated in technical triplicate in wells containing the test compound and fresh ATc (100 ng/mL). Δ*rv3645* strains were cultured in 7H9-ADC prior to back diluting in 7H9-OADC to seeding a 384-well plate. Similarly, for strains used in antibacterial activity testing in the presence or absence of fatty acids, strains were grown in 7H9-ADC before back diluting the culture to a starting OD_600_ of 0.05 in 7H9-ADC or 7H9-OADC to seed a 384-well plate. Plates were incubated at 37°C with 5% CO_2_. OD_600_ was evaluated using a Tecan Spark plate reader at 14-18 days post-plating and percent growth was calculated relative to the vehicle control for each strain. IC_50_ measurements were calculated using a non-linear fit in GraphPad Prism.

For fatty acid sensitivity measurements, fatty acids were dissolved in 1:1 tyloxapol:ethanol and then diluted at 2X the maximum testing concentration in 7H9-ADC. 2-fold serial dilutions were prepared and 25 μL of each concentration was transferred to a 384-well plate. Cultures were grown to OD_600_ 0.4-0.6. Cultures were then back diluted to a starting OD_600_ of 0.1 and 25 μL of cell suspension was plated in technical triplicate in wells containing 25 μL of the fatty acid dilution series (and fresh ATc (100 ng/mL), where applicable).

To quantify growth phenotypes on 7H10 agar, 10-fold serial dilutions of OD-synchronized Mtb cultures were spotted on 7H10-ADC agar containing fatty acids at the indicated concentrations. Where applicable, ATc was added at 100 ng/mL. Plates were incubated at 37°C and imaged after two weeks.

### CELL WALL PERMEABILITY ASSAY

Cell envelope permeability was determined using the ethidium bromide (EtBr) uptake assay as previously described (Xu et al., 2017). Briefly, mid-log-phase Mtb cultures were washed once in PBS + 0.05% Tween-80 and adjusted to OD_600_=0.8 in PBS supplemented with 0.4% glucose. 100 μL of bacterial suspension was added to a black 96-well clear-bottomed plate (Costar). After this, 100 μL of 8 μg/mL EtBr in PBS supplemented with 0.4% glucose was added to each well. EtBr fluorescence was measured (excitation: 530 nm/emission: 590 nm) at 1 min intervals over a course of 90 min. EtBr fluorescence at 30 min is plotted, normalized to optical density. Experiments were performed in technical sextuplicate.

A similar assay was performed to determine envelope permeability to a fluorescent vancomycin analogue, except that: (1) the bacterial suspension was adjusted at OD_600_=0.4 in PBS supplemented with 0.4% glucose; (2) cells were incubated with 2 μg/mL BODIPY FL Vancomycin (Thermo Scientific, V34850) for 30 min; (3) 900 μL sample aliquots were taken at different time points, washed twice with PBS, resuspended in 600 μL PBS, and 3 aliquots of 200 uL each were transferred to a black 96-well clear-bottomed plate (Costar); and (4) fluorescence was measured (excitation: 485 nm/emission: 538 nm) and normalized to the OD_600_ of the final bacterial suspension.

### MEASURING DRUG UPTAKE BY MASS-SPECTROMETRY

CRISPRi strains were growth-synchronized and pre-depleted in the presence of ATc (100 ng/mL) for 4 days. Growth in 7H9-OADC ATc media proceeded to an OD of 0.8-1.0. 1 OD unit was used to seed filters for growth on 7H10-OADC ATc agar plates. After 5 days of growth, bacteria-laden filters were transferred and floated on 7H9-OADC ATc media in a “swimming pool” set up overnight. Filters were then transferred and floated on drug-containing 7H9-OADC ATc swimming pools and incubated for 24 hours. Media was collected from each pool and filter sterilized using 0.22-micron nylon centrifugal filters (Corning). 100 μL of each media sample was added to 400 μL of 1:1 acetonitrile/methanol and centrifuged at 13,000 *g* for 10 minutes at 4°C to pellet precipitated protein. Supernatants were transferred into mass spectrometry vials and analyzed using a semi-quantitative LC/MS-based method as described previously (Planck and Rhee, 2021). Briefly, samples were separated on a Cogent Diamond Hydride Type C column (Microsolv Technologies). The mobile phase consisted of solvent A (ddH_2_O with 0.2% formic acid) and solvent B (acetonitrile with 0.2% formic acid), and the gradient used was as follows: 0–2 min, 85% B; 3–5 min, 80% B; 6–7 min, 75% B; 8–9 min, 70% B; 10–11.1 min, 50% B; 11.1–14 min 20% B; 14.1–24 min 5% B, followed by a 10 min re-equilibration period at 85% B at a flow rate of 0.4 mL/min. This was achieved using an Agilent 1200 Series liquid chromatography (LC) system coupled to an Agilent 6546 quadrupole time of flight (Q-TOF) mass spectrometer in positive acquisition mode, and 2 μL of sample were injected for each run. Dynamic mass axis calibration was achieved by continuous infusion of a reference mass solution using an isocratic pump with a 100:1 splitter. Resulting data were analyzed using Agilent MassHunter Qualitative Analysis Navigator software. Relative antibiotic concentrations were determined by quantitation of peak heights using *m/z* of 338.1511 for the linezolid (M+H)^+^ ion (RT = 1.3 min) and 724.7224 for the vancomycin (M+2H)^2+^ ion (RT = 8.8 min) with a mass tolerance of +/-30 ppm. To determine uptake, drug levels in bacteria-laden filter swimming pools were compared to levels from control swimming pools incubated with cell-free filters under the same conditions.

### GENERATION OF INDIVIDUAL CRISPRI AND CRISPRI-RESISTANT COMPLEMENTATION STRAINS

Individual CRISPRi plasmids were cloned as previously described in (Bosch et al., 2021) using Addgene plasmid #166886. Briefly, the CRISPRi plasmid backbone was digested with BsmBI-v2 (NEB #R0739L) and gel purified. sgRNAs were designed to target the non-template strand of the target gene ORF. For each individual sgRNA, two complementary oligonucleotides with appropriate sticky end overhangs were annealed and ligated (T4 ligase NEB # M0202M) into the BsmBI-digested plasmid backbone. Successful cloning was confirmed by Sanger sequencing.

Individual CRISPRi plasmids were then electroporated into Mtb. Electrocompetent cells were obtained as described in (Murphy et al., 2015). Briefly, a WT Mtb culture was expanded to an OD_600_=0.8-1.0 and pelleted (4,000 x g for 10 min). The cell pellet was washed three times in sterile 10% glycerol. The washed bacilli were then resuspended in 10% glycerol in a final volume of 5% of the original culture volume. For each transformation, 100 ng plasmid DNA and 100 μL of electrocompetent mycobacteria were mixed and transferred to a 2 mm electroporation cuvette (Bio-Rad #1652082). Where necessary, 100 ng of plasmid plRL19 (Addgene plasmid #163634) was also added. Electroporation was performed using the Gene Pulser X cell electroporation system (Bio-Rad #1652660) set at 2500 V, 700 Ω and 25 μF. Bacteria were recovered in 7H9 for 24 hours. After the recovery incubation, cells were plated on 7H10 agar supplemented with the appropriate antibiotic to select for transformants.

To complement CRISPRi-mediated gene knockdown, synonymous mutations were introduced into the complementing allele at both the protospacer adjacent motif (PAM) and seed sequence (the 8-10 most PAM-proximal bases at the 3’ end of the sgRNA targeting sequence) to prevent sgRNA targeting, as described here (Wong and Rock, 2021). Silent mutations were introduced into Gibson assembly oligos to generate these “CRISPRi resistant” (CR) alleles. Complementation alleles were expressed from hsp60 promoters in a Tweety integrating plasmid backbone, as indicated in each figure legend and/or the relevant plasmid maps (**Supplemental Table 1**). These alleles were then transformed into the corresponding CRISPRi knockdown strain.

The full list of sgRNA targeting sequences and complementation plasmids can be found in **Supplemental Table 1**.

### CONSTRUCTION OF THE Δ*RV3645* AND COMPLEMENTED MTB STRAINS

Mtb H37Rv gene *rv3645* was deleted through homologous recombination using a strain expressing the recombinase RecET (Murphy et al., 2015). To induce RecET expression, isovaleronitrile was added at a final concentration of 1 μM to a mid-exponential Mtb culture (optical density 580nm of approximately 1) for 8 hours after which glycine was added at a final concentration of 2M and the culture was incubated overnight. A construct composed of a hygromycin resistant gene (hygR) flanked by 500 bp upstream and downstream of *rv3645* was synthesized (GenScript) and electroporated into Mtb expressing the recombinase RecET. A *rv3645* deletion mutant strain (Δ*rv3645*) was selected in solid fatty acid free modified Sauton’s with hygromycin. The plasmid expressing recET (pNitET-SacB-kan) was counterselected by growing the deleted mutant in solid fatty acid free modified Sauton’s supplemented with sucrose 10%. For complementation of Δ*rv3645*, we have cloned *rv3645* under the control of the promoter Phsp60 into a plasmid with a kanamycin resistant cassette that integrates at the att-L5 site (pMCK-Phsp60-rv3645) and electroporated it into rv3645 deletion mutant. Primers, plasmids and strains are listed in Supplementary Tables 3 and 4, respectively.

### ISOLATION OF SPONTANEOUS FATTY ACID TOXICITY SUPPRESSORS

The mutant strain Δ*rv3645* was grown in fatty acid free modified Sauton’s minimal medium until stationary phase (Beites et al., 2019). Solid fatty acid free modified Sauton’s minimal medium supplemented with oleic acid at a final concentration of 500 μM was used to select spontaneous mutants in the Δ*rv3645* genetic background that regained the ability to grow in the presence of oleic acid. We inoculated this medium with 10^7^ and 10^8^ bacteria and incubated the plates for 4 weeks. Medium not supplemented with oleic acid was used as a viability control. Colonies that grew in the medium supplemented with oleic acid were picked and grown in liquid fatty acid free modified Sauton’s minimal medium. To validate the isolated rescue mutants, we cultured WT, Δ*rv3645*, complemented and rescue mutants in liquid fatty acid free modified Sauton’s minimal medium supplemented with a concentration of oleic acid restrictive to Δ*rv3645* growth.

### WHOLE GENOME SEQUENCING

The genetic identity of Δ*rv3645* and derived spontaneous suppressor mutants was confirmed by whole genome sequencing (WGS). Genomic DNA (150-200 ng) was sheared and HiSeq sequencing libraries were prepared using the KAPA Hyper Prep Kit (Roche). Libraries were amplified by PCR (10 cycles). 5– 10 × 10^6^ 50-bp paired-end reads were obtained for each sample on an Illumina HiSeq 2500 using the TruSeq SBS Kit v3 (Illumina). Post-run demultiplexing and adapter removal were performed and FASTQ files were inspected using fastqc (Andrews, 2010). Trimmed FASTQ files and the reference genome (M. tuberculosis H37RvCO; NZ_CM001515.1) were aligned using bwa mem (Li and Durbin, 2010). Bam files were sorted and merged using SAMtools (Li et al., 2009). Read groups were added and bam files de-duplicated using Picard tools and GATK best-practices were followed for SNP and indel detection (DePristo et al., 2011). Gene knockouts and cassette insertions were verified for all strains by direct comparison of reads spanning insertion points to plasmid maps and the genome sequence. Reads coverage data was obtained from the software Integrative Genomics Viewer version 2.5.2 (IGV) (Robinson et al., 2011).

### CONSTRUCTION OF A GENOME-WIDE CRISPRI LIBRARY IN Δ*RV3645*

Libraries were constructed as previously described (Bosch et al., 2021). Briefly thirty seven transformations were performed to generate Δ*rv3645* RLC12 libraries. For each transformation, 1 μg of RLC12 plasmid DNA was added to 100 μL electrocompetent cells (∼1 × 10^10^ cells per transformation). The cells:DNA mix was transferred to a 2 mm electroporation cuvette (Bio-Rad #1652082) and electroporated at 2500 kV, 700 ohms, and 25 μF. Each transformation was recovered in 2 mL Sauton’s media supplemented with fatty acid-free ADC, glycerol and tyloxapol (80 mL total) for 16-24 hours. The recovered cells were harvested at 4,000 rpm for 10 minutes, resuspended in 700 μL remaining media per transformation and plated on Sauton’s agar supplemented with kanamycin (see Bacterial cultures) in Corning Bioassay dishes (Sigma #CLS431111-16EA). Transformation efficiency was estimated from library titring and indicated > 12X average sgRNA coverage of RLC12 was achieved in Δ*rv3645*.

After 33 days of outgrowth on plates, transformants were scraped and pooled. Scraped cells were homogenized by two dissociation cycles on a gentleMACS Octo Dissociator (Miltenyi Biotec #130095937) using the RNA_02.01 program and 10 gentleMACS M tubes (Miltenyi Biotec #130093236). The library was further declumped by passaging 10 individual *M. tuberculosis* library aliquots in 10 mL of kanamycin supplemented Sauton’s in T-25 flasks (Falcon # 08-772-1F) for 10 generations. Final Δ*rv3645* RLC12 library stocks were obtained after pooling the cultures and passing them through a 10 μm cell strainer (Pluriselect #SKU 43-50010-03). Genomic DNA was extracted from the final Δ*rv3645* RLC12 library stock and library quality was validated by deep sequencing (see Genomic DNA extraction and library preparation for Illumina sequencing).

### CRISPRI FATTY ACID-GENETIC SUPPRESSOR SCREENING

The fatty acid-genetic suppressor screen was initiated by thawing 3 × 1.5 mL aliquots (1 OD_600_ unit per aliquot) of the **Δ***rv3645* CRISPRi library and inoculating each aliquot into 8.5 mL 7H9-ADC in a vented tissue culture flask (T-25; Corning #430639). The starting OD_600_ of each culture was approximately 0.1. Cultures were expanded to OD_600_=0.47, pooled, and evenly divided to inoculate 2 × 90 mL cultures with 7.5 ODU each in tissue culture flasks (T-225; Falcon #353138). Cultures were expanded to OD 0.3, pooled, pelleted, and resuspended in 15 mL 7H9-ADC. 700 μL of the concentrated cells were plated on FA-free 7H10-ADC 25 cm bioassay dishes, or 7H10-ADC with increasing concentrations of palmitic acid (200 μM) in quintuplicate. Bioassay dishes were supplemented with kanamycin at 20 μg/mL and ATc at 100 ng/mL. To titer the library, a 10-fold dilution series of the concentrated cells was plated on petri dishes with FA-free 7H10-ADC with kanamycin at 20 μg/mL. All plates were incubated for 20 days. Library coverage based on titering plates was 4620X. Colonies from the fatty acid-containing bioassay dishes were scraped, avoiding clustered colonies, into PBS and pelleted. Due to confluent growth in the absence of selection on the fatty acid-free plates, a 3 cm x 25 cm rectangular area was scraped into PBS and cells were pelleted for genomic DNA extraction.

## DATA AVAILABILITY

Whole genome sequencing data for Δ*rv3645* and derived spontaneous rescue mutants were deposited in NCBI’s Sequence Read Archive (SRA) under BioProject PRJNA811534. CRISPRi suppressor screen sequencing data was deposited in NCBI’s SRA under BioProject PRJNA814682.

### GENOMIC DNA EXTRACTION AND LIBRARY PREPARATION FOR ILLUMINA SEQUENCING

Genomic DNA was isolated from bacterial pellets using the CTAB-lysozyme method as previously described as previously described (Bosch et al., 2021) Larsen et al. (Larsen et al., 2007). Genomic DNA concentration was quantified by Nanodrop. Next, the sgRNA-encoding region was amplified from 500 ng of genomic DNA using NEBNext Ultra II Q5 master Mix (NEB #M0544L). PCR cycling conditions were: 98°C for 45 s; 17 cycles of 98°C for 10 s, 64°C for 30 s, 65°C for 20 s; 65°C for 5 min. Samples were dual-indexed. For dual-indexed samples, each PCR reaction contained a unique indexed forward and reverse primer (0.5 μM each) (**Table S1**). Forward primers contain a P5 flow cell attachment sequence, a standard Read1 Illumina sequencing primer binding site, custom stagger sequences to ensure base diversity during Illumina sequencing, and a unique barcode to allow for sample pooling during deep sequencing. Reverse primers contain a P7 flow cell attachment sequence, a standard Read2 Illumina sequencing primer binding site, and unique barcodes to allow for dual-indexed sequencing.

Following PCR amplification, each ∼230 bp amplicon was purified using using sparQ PureMag Beads (Quantabio # 95196-060) using double-sided size selection (first 0.75X, then an additional 0.12X for a final 0.87X). Size-selected amplicons were quantified with a Qubit 2.0 Fluorometer (Invitrogen). Amplicon size and purity were quality controlled by visualization on an Agilent 2100 Bioanalyzer (high sensitivity chip; Agilent Technologies #5067-4626). Next, individual PCR amplicons were multiplexed into 10 nM pools and sequenced on an Illumina sequencer according to the manufacturer’s instructions (2.5-5% PhiX spike-in; PhiX Sequencing Control v3; Illumina # FC-110-3001). Samples were run on the Illumina NextSeq 500 platform (Single-Read 1×85 cycles and six i7 index cycles).

### WESTERN BLOTTING

80 OD_600_ units of growth synchronized *M. smegmatis* cultures were harvested by centrifugation (4,000 x g, 10 min). Cells were washed twice in 40 mL PBS-0.05% Tween80 and resuspended in 600 μL of lysis buffer (50 mM Tris, 150 mM NaCl, pH 7.4) containing a protease inhibitor cocktail (Sigma-Aldrich, #11873580001). Cells were lysed by bead beating in Lysis B Matrix tubes (MP Biomedicals; #116911050) using a Precellys Evolution homogenizer (Bertin Instruments, #P000062-PEVO0-A, 3 × 10,000 RPM, 30 s intervals, 4°C). n-Dodecyl-β-D-maltopyranoside (Alfa Aesar, #J66869) was added to a final concentration of 1% and incubated at 4°C with inversion for 2 h. The cell lysates were cleared by centrifugation (20,000 x g, 2 min) and a 20 μL aliquot was mixed with 4X Laemmli Sample Buffer (Bio-Rad, #1610747) supplemented with DTT. Samples were separated on a 4-12% Bis-Tris polyacrylamide gel (Invitrogen, #NP0323BOX) in MOPS running buffer, transferred to a nitrocellulose membrane using the TransBlot Turbo Transfer System (Bio-Rad, #1704150), and incubated for 1 hour in blocking buffer (LI-COR, #927-60001). Proteins were probed with anti-RpoB (BioLegend, #663905) and anti-His (GenScript, #A00186) primary antibodies overnight at 4°C and subsequently detected with fluorescent goat anti-mouse secondary antibodies (Bio-Rad, #12004159).

### MEASURING CAMP BY MASS SPECTROMETRY

Strains were grown to an OD of 0.8-1.0 in 7H9-ADC media. CRISPRi strains were grown for 4 days to an OD of 0.8-1.0 in the presence of ATc (100 ng/mL) to predeplete targets. For CRISPRi strains, ATc was maintained in the media at the same concentration until cells were harvested. 1 OD unit was used to seed filters for growth on 7H10-ADC agar plates. After 5 days of growth, bacteria-laden filters were transferred and floated on 7H9-ADC media in a “swimming pool” set up overnight. Filters were then transferred and floated on 7H9-OADC swimming pools and incubated for 24 hours. For metabolite extraction, filters were transferred to 1 mL of acetonitrile:methanol:water (2:2:1). Bacteria were disrupted by bead beating 6 times at 6,000 rpm for 30 s at 4°C (Precellys), and lysates were clarified by centrifugation and filter sterilized as described above. Lysates were transferred into mass spectrometry vials and analyzed using a semi-quantitative LC/MS-based ion pairing method. Briefly, samples were separated on a Zorbax Extend C18 column (Agilent). The mobile phase consisted of solvent A (97:3 water:methanol) and solvent B (100% methanol), both containing 5 mM tributylamine and 5.5 mM acetic acid, and the gradient used was as follows: 0–3.5 min, 0% B; 4–7.5 min, 30% B; 8–15 min, 35% B; 20–24 min, 99% B; 24–24.5 min, 0% B, followed by a 10 min re-equilibration period at 0% B at a flow rate of 0.25 mL/min. This was achieved using an Agilent 1200 Series liquid chromatography (LC) system coupled to an Agilent 6220 accurate mass time of flight (TOF) mass spectrometer in negative acquisition mode, and 5 μL of sample were injected for each run. Dynamic mass axis calibration was achieved by continuous infusion of a reference mass solution using an isocratic pump with a 100:1 splitter. Resulting data were analyzed using Agilent MassHunter Qualitative Analysis Navigator software. Relative abundances of cAMP were determined by quantitation of peak heights using *m/z* of 328.0452 for the cAMP (M-H)^-^ ion (RT = 9.0 min) with a mass tolerance of +/-35 ppm. Absolute abundances of cAMP were determined by comparing sample peak heights to a standard curve generated by spiking cAMP (final concentration range 0.015 μM to 3.9 μM) into mycobacterial lysate. In all cases, ion counts were normalized to residual protein content in the samples, which was measured using a BCA assay (Pierce).

## SUPPLEMENTAL INFORMATION

**Supplemental Table 1:**
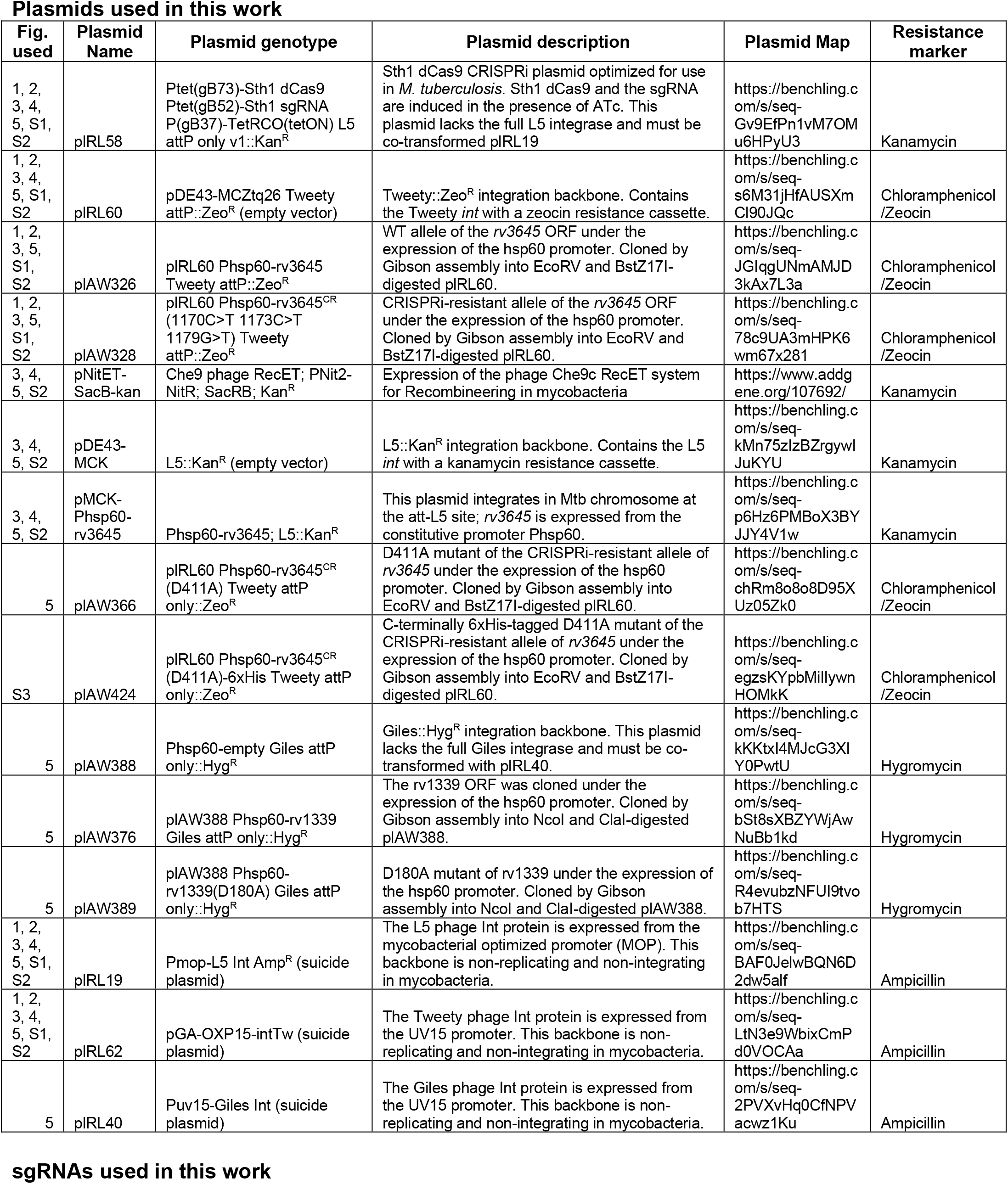
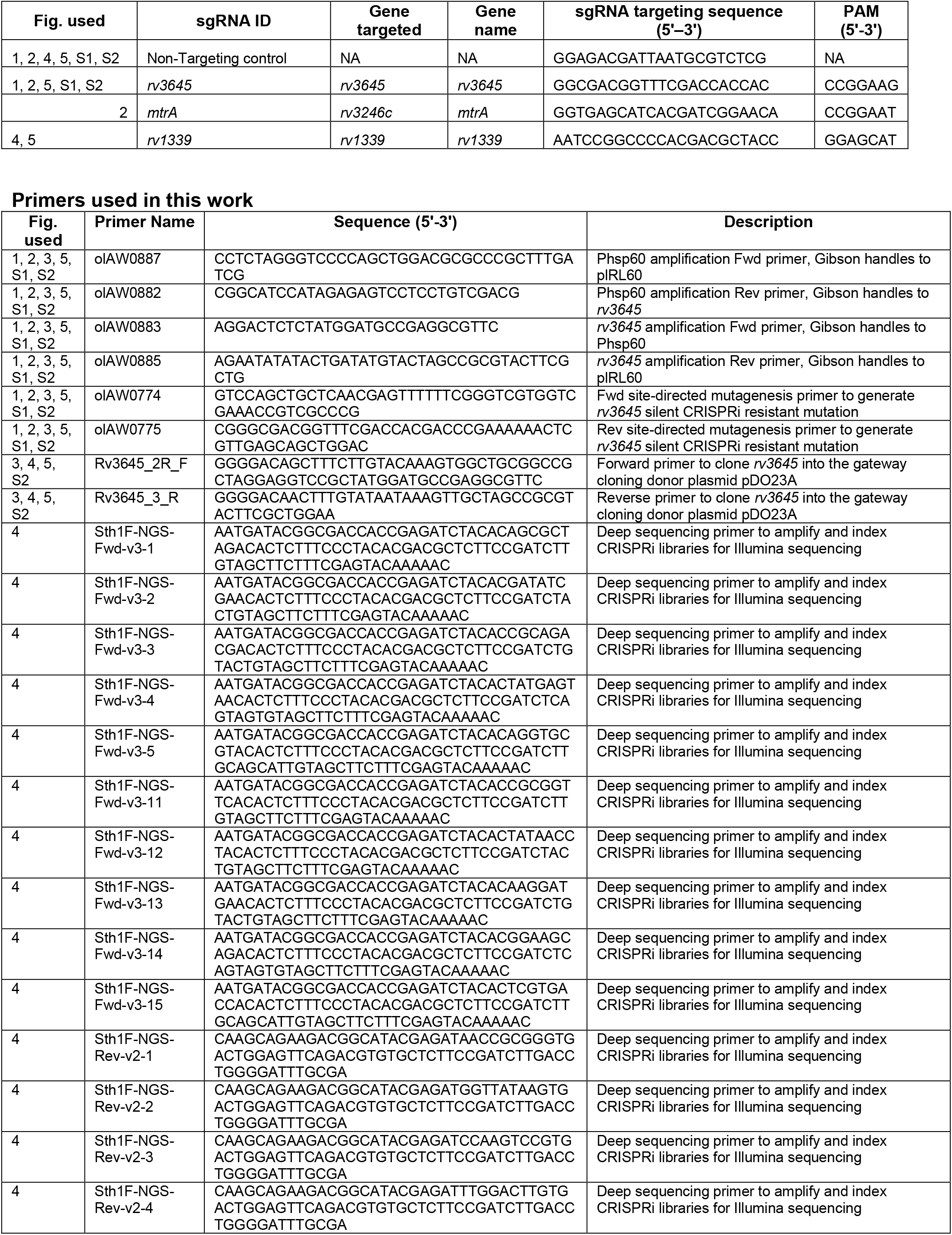
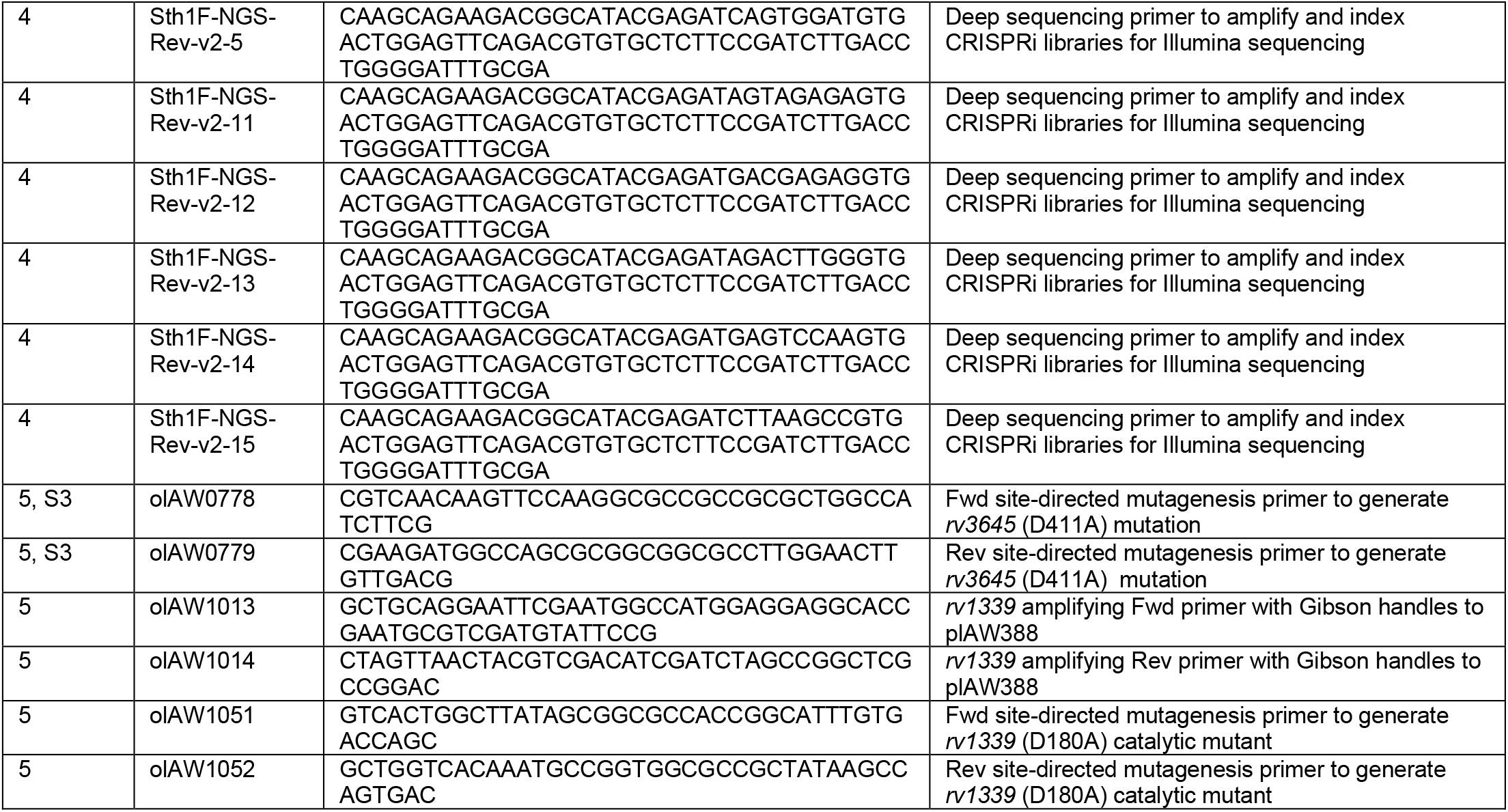
List of plasmids and primers used in this work

## SUPPLEMENTAL FIGURES

**Supplemental Figure 1:**
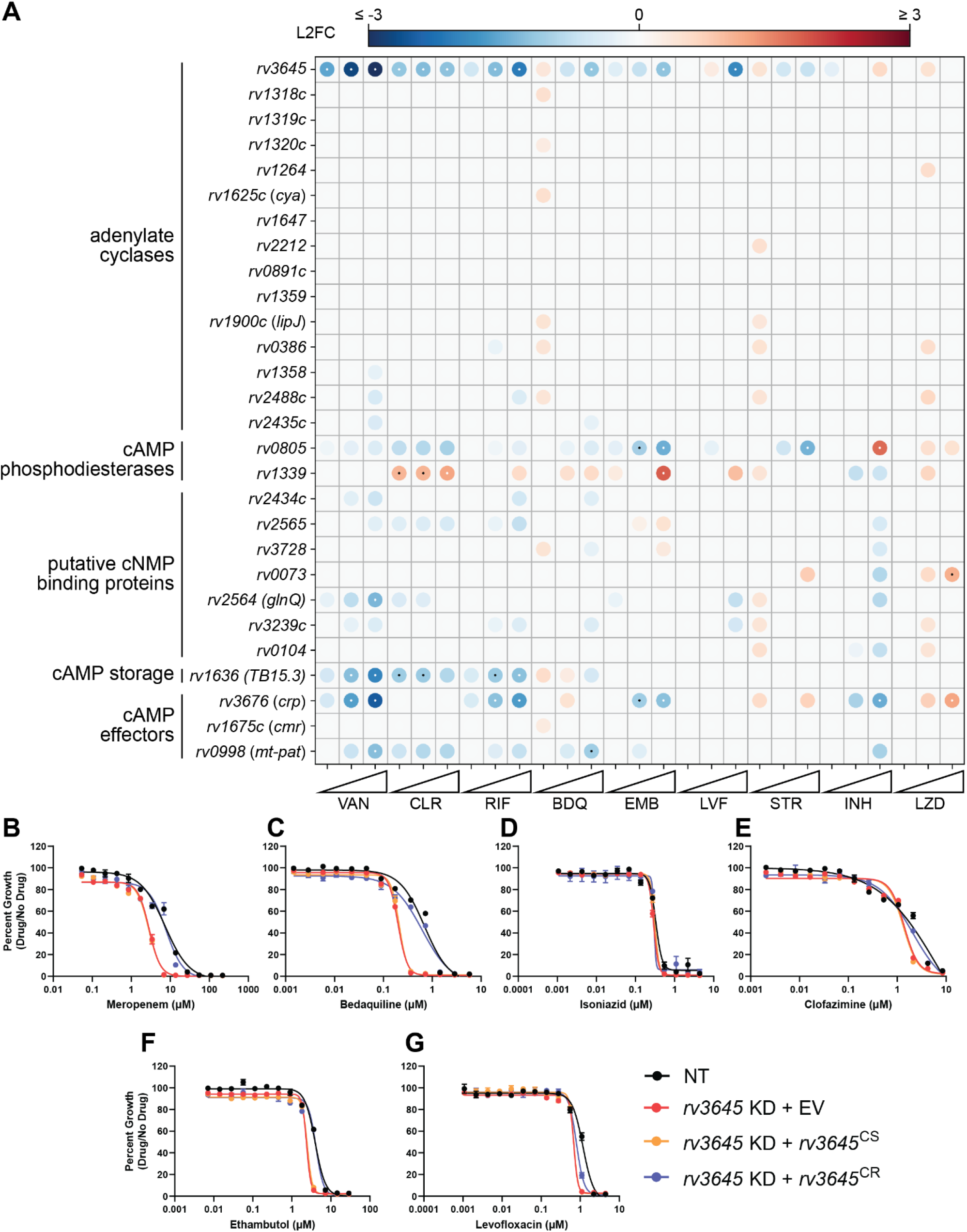
In chemical-genetic CRISPRi screens, *rv3645* is a unique hit among other genes encoding adenylate cyclases, cAMP phosphodiesterases, and cAMP effectors. (A) Feature-expression heatmap of annotated adenylate cyclase, cAMP phosphodiesterase, putative cNMP binding protein, cAMP storage, and cAMP effector genes from a 5-day CRISPRi library pre-depletion screen (Li et al., 2022). The color of each circle represents the gene-level L2FC; a white dot represents an FDR < 0.01 and a |L2FC| > 1. VAN = vancomycin; CLR = clarithromycin; RIF = rifampicin; BDQ = bedaquiline; EMB = ethambutol; LVX = levofloxacin; STR = streptomycin; INH = isoniazid; LZD = linezolid. (B-G) Dose-response curves for meropenem (B), bedaquiline (C), isoniazid (D), clofazimine (E), ethambutol (F), and levofloxacin (G). NT = non-targeting sgRNA; KD = knockdown; CS = CRISPRi-sensitive; CR = CRISPRi-resistant. Data represent mean ± SEM for technical triplicates and are representative of three independent experiments.

**Supplemental Figure 2:**
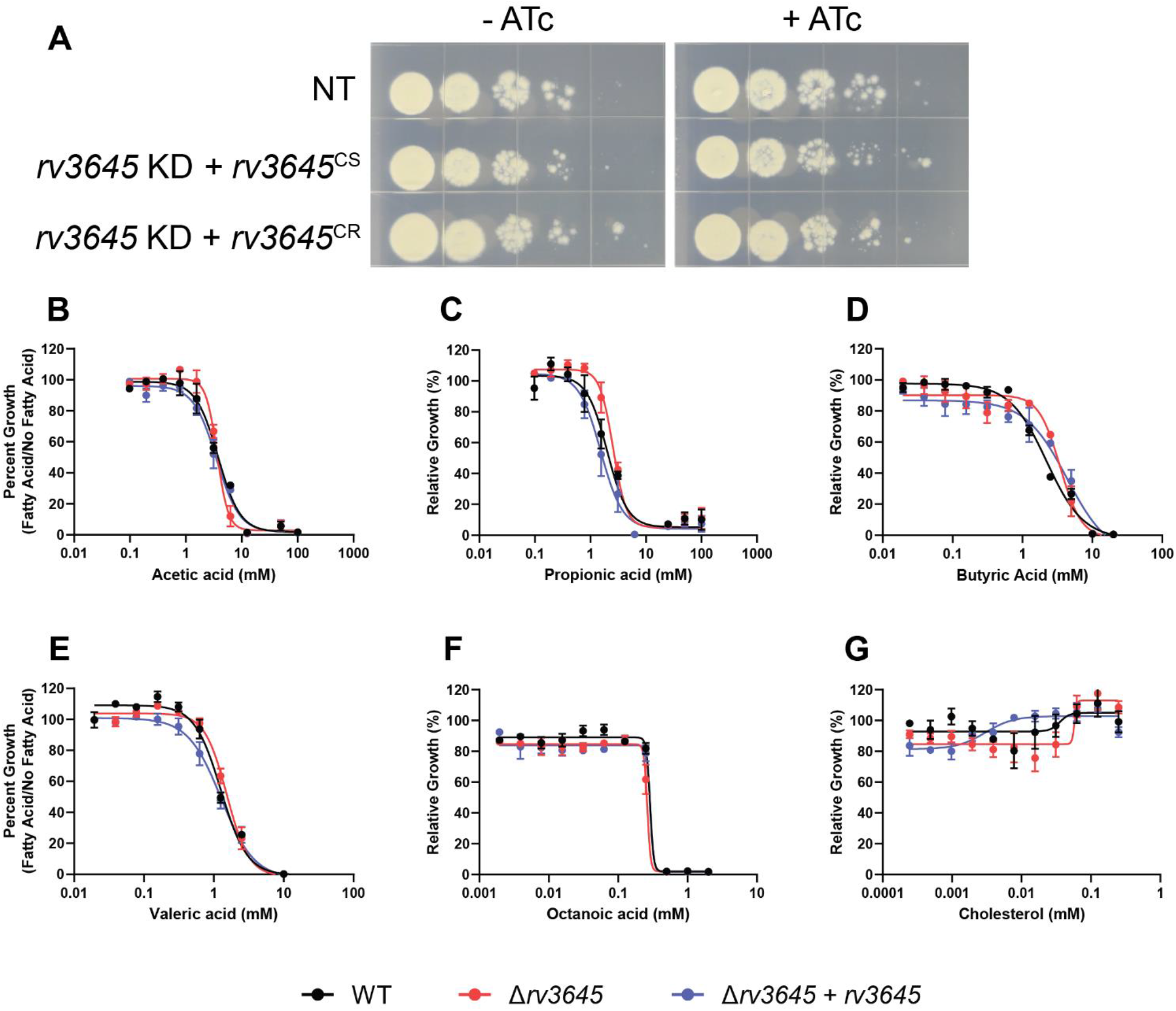
Deletion of *rv3645* does not sensitize Mtb to short-or medium-chain fatty acids or cholesterol. (A) Growth of *rv3645* CRISPRi strains on fatty acid-free 7H10-ADC agar. NT = non-targeting sgRNA; KD = knockdown; CS = CRISPRi-sensitive; CR = CRISPRi-resistant. (B-G) Dose-response curves for short-chain fatty acids acetic acid (B), propionic acid (C), butyric acid (D), and valeric acid (E); medium-chain fatty acid octanoic acid (F), and cholesterol (G). Data represent mean ± SEM for technical triplicates and are representative of three independent experiments.

**Supplemental Figure 3:**
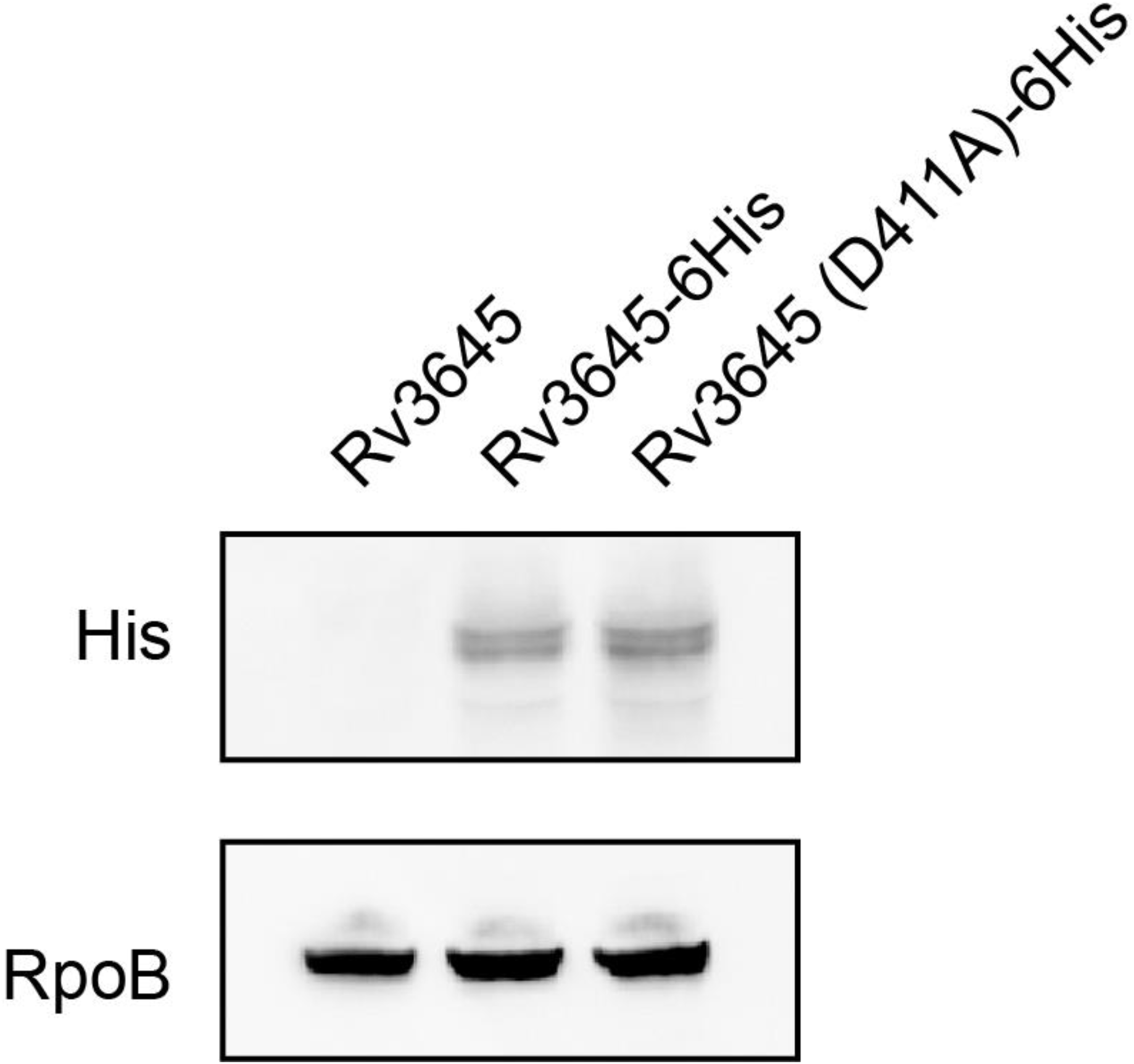
*rv3645* catalytic mutant is expressed in mycobacteria. Western blot for C-terminally 6His-tagged *rv3645* alleles expressed in *M. smegmatis*. Western blot for RpoB as loading control. Note the double band present in the anti-His blot may reflect processing of the predicted N-terminal signal peptide of Rv3645.

**Supplemental Figure 3 – source data 1** Original Western blot image of *rv3645* catalytic mutant.

## REFERENCES

Abdel Motaal A, Tews I, Schultz JE, Linder JU. 2006. Fatty acid regulation of adenylyl cyclase Rv2212 from Mycobacterium tuberculosis H37Rv. FEBS J 273:4219–28. doi:10.1111/j.1742-4658.2006.05420.x

Agarwal N, Lamichhane G, Gupta R, Nolan S, Bishai WR. 2009. Cyclic AMP intoxication of macrophages by a Mycobacterium tuberculosis adenylate cyclase. Nature 460:98–102. doi:10.1038/nature08123

Andrews S. 2010. FastQC: A Quality Control Tool for High Throughput Sequence Data. http://www.bioinformatics.babraham.ac.uk/projects/fastqc/

Bai G, Schaak DD, McDonough KA. 2009. cAMP levels within Mycobacterium tuberculosis and Mycobacterium bovis BCG increase upon infection of macrophages. FEMS Immunol Med Microbiol 55:68–73. doi:10.1111/j.1574-695X.2008.00500.x

Batt SM, Minnikin DE, Besra GS. 2020. The thick waxy coat of mycobacteria, a protective layer against antibiotics and the host’s immune system. Biochem J 477:1983. doi:10.1042/BCJ20200194

Beites T, O’Brien K, Tiwari D, Engelhart CA, Walters S, Andrews J, Yang H-J, Sutphen ML, Weiner DM, Dayao EK, Zimmerman M, Prideaux B, Desai P V., Masquelin T, Via LE, Dartois V, Boshoff HI, Barry CE, Ehrt S, Schnappinger D. 2019. Plasticity of the Mycobacterium tuberculosis respiratory chain and its impact on tuberculosis drug development. Nat Commun 10:4970. doi:10.1038/s41467-019-12956-2

Bellerose MM, Proulx MK, Smith CM, Baker RE, Ioerger TR, Sassetti CM. 2020. Distinct Bacterial Pathways Influence the Efficacy of Antibiotics against Mycobacterium tuberculosis. mSystems 5. doi:10.1128/mSystems.00396-20

Bosch B, DeJesus MA, Poulton NC, Zhang W, Engelhart CA, Zaveri A, Lavalette S, Ruecker N, Trujillo C, Wallach JB, Li S, Ehrt S, Chait BT, Schnappinger D, Rock JM. 2021. Genome-wide gene expression tuning reveals diverse vulnerabilities of M. tuberculosis. Cell 184(17):4579–4592.

Boutte CC, Baer CE, Papavinasasundaram K, Liu W, Chase MR, Meniche X, Fortune SM, Sassetti CM, Ioerger TR, Rubin EJ. 2016. A cytoplasmic peptidoglycan amidase homologue controls mycobacterial cell wall synthesis. Elife 5. doi:10.7554/ELIFE.14590

Cann MJ, Hammer A, Zhou J, Kanacher T. 2003. A Defined Subset of Adenylyl Cyclases Is Regulated by Bicarbonate Ion. J Biol Chem 278:35033–35038. doi:10.1074/JBC.M303025200

Chandrasekera NS, Berube BJ, Shetye G, Chettiar S, O’Malley T, Manning A, Flint L, Awasthi D, Ioerger TR, Sacchettini J, Masquelin T, Hipskind PA, Odingo J, Parish T. 2017. Improved Phenoxyalkylbenzimidazoles with Activity against Mycobacterium tuberculosis Appear to Target QcrB. ACS Infect Dis 3:898–916. doi:10.1021/acsinfecdis.7b00112

DeJesus MA, Gerrick ER, Xu W, Park SW, Long JE, Boutte CC, Rubin EJ, Schnappinger D, Ehrt S, Fortune SM, Sassetti CM, Ioerger TR. 2017. Comprehensive Essentiality Analysis of the Mycobacterium tuberculosis Genome via Saturating Transposon Mutagenesis. MBio 8. doi:10.1128/mBio.02133-16

DePristo MA, Banks E, Poplin R, Garimella K V, Maguire JR, Hartl C, Philippakis AA, del Angel G, Rivas MA, Hanna M, McKenna A, Fennell TJ, Kernytsky AM, Sivachenko AY, Cibulskis K, Gabriel SB, Altshuler D, Daly MJ. 2011. A framework for variation discovery and genotyping using next-generation DNA sequencing data. Nat Genet 43:491–8. doi:10.1038/ng.806

Eoh H, Rhee KY. 2014. Methylcitrate cycle defines the bactericidal essentiality of isocitrate lyase for survival of mycobacterium tuberculosis on fatty acids. Proc Natl Acad Sci U S A 111:4976–4981. doi:10.1073/pnas.1400390111

Eoh H, Rhee KY. 2013. Multifunctional essentiality of succinate metabolism in adaptation to hypoxia in Mycobacterium tuberculosis. Proc Natl Acad Sci U S A 110:6554–6559. doi:10.1073/PNAS.1219375110

Galagan JE, Minch K, Peterson M, Lyubetskaya A, Azizi E, Sweet L, Gomes A, Rustad T, Dolganov G, Glotova I, Abeel T, Mahwinney C, Kennedy AD, Allard R, Brabant W, Krueger A, Jaini S, Honda B, Yu WH, Hickey MJ, Zucker J, Garay C, Weiner B, Sisk P, Stolte C, Winkler JK, Van De Peer Y, Iazzetti P, Camacho D, Dreyfuss J, Liu Y, Dorhoi A, Mollenkopf HJ, Drogaris P, Lamontagne J, Zhou Y, Piquenot J, Park ST, Raman S, Kaufmann SHE, Mohney RP, Chelsky D, Branch Moody D, Sherman DR, Schoolnik GK. 2013. The Mycobacterium tuberculosis regulatory network and hypoxia. Nature 499:178–183. doi:10.1038/NATURE12337

Gazdik MA, McDonough KA. 2005. Identification of cyclic AMP-regulated genes in Mycobacterium tuberculosis complex bacteria under low-oxygen conditions. J Bacteriol 187:2681–92. doi:10.1128/JB.187.8.2681-2692.2005

Hulko M, Berndt F, Gruber M, Linder JU, Truffault V, Schultz A, Martin J, Schultz JE, Lupas AN, Coles M. 2006. The HAMP domain structure implies helix rotation in transmembrane signaling. Cell 126:929–40. doi:10.1016/j.cell.2006.06.058

Johnson Richard M, McDonough KA. 2018. Cyclic nucleotide signaling in Mycobacterium tuberculosis: an expanding repertoire. Pathog Dis 76. doi:10.1093/femspd/fty048

Johnson Richard M., McDonough KA. 2018. Cyclic nucleotide signaling in Mycobacterium tuberculosis: an expanding repertoire. Pathog Dis 76. doi:10.1093/femspd/fty048

Kengmo Tchoupa A, Eijkelkamp BA, Peschel A. 2022. Bacterial adaptation strategies to host-derived fatty acids. Trends Microbiol 30:241–253. doi:10.1016/j.tim.2021.06.002

Ko EM, Oh J Il. 2020. Induction of the cydAB Operon Encoding the bd Quinol Oxidase Under Respiration-Inhibitory Conditions by the Major cAMP Receptor Protein MSMEG_6189 in Mycobacterium smegmatis. Front Microbiol 11. doi:10.3389/FMICB.2020.608624

Larrouy-Maumus G, Marino LB, Madduri AVR, Ragan TJ, Hunt DM, Bassano L, Gutierrez MG, Moody DB, Pavan FR, de Carvalho LPS. 2016. Cell-Envelope Remodeling as a Determinant of Phenotypic Antibacterial Tolerance in Mycobacterium tuberculosis. ACS Infect Dis 2:352–360. doi:10.1021/acsinfecdis.5b00148

Larsen MH, Biermann K, Tandberg S, Hsu T, Jacobs WRJ. 2007. Genetic Manipulation of Mycobacterium tuberculosis. Curr Protoc Microbiol Chapter 10:Unit 10A.2. doi:10.1002/9780471729259.mc10a02s6

Li H, Durbin R. 2010. Fast and accurate long-read alignment with Burrows-Wheeler transform. Bioinformatics 26:589–95. doi:10.1093/bioinformatics/btp698

Li H, Handsaker B, Wysoker A, Fennell T, Ruan J, Homer N, Marth G, Abecasis G, Durbin R, 1000 Genome Project Data Processing Subgroup. 2009. The Sequence Alignment/Map format and SAMtools. Bioinformatics 25:2078–9. doi:10.1093/bioinformatics/btp352

Li S, Poulton NC, Chang JS, Azadian ZA, DeJesus MA, Ruecker N, Zimmerman MD, Eckartt KA, Bosch B, Engelhart CA, Sullivan DF, Gengenbacher M, Dartois VA, Schnappinger D, Rock JM. 2022. CRISPRi chemical genetics and comparative genomics identify genes mediating drug potency in Mycobacterium tuberculosis. Nat Microbiol 7:766–779. doi:10.1038/s41564-022-01130-y

Li W, Xu H, Xiao T, Cong L, Love MI, Zhang F, Irizarry RA, Liu JS, Brown M, Liu XS. 2014. MAGeCK enables robust identification of essential genes from genome-scale CRISPR/Cas9 knockout screens. Genome Biol 15:554. doi:10.1186/S13059-014-0554-4

Linder JU, Schultz A, Schultz JE. 2002. Adenylyl cyclase Rv1264 from Mycobacterium tuberculosis has an autoinhibitory N-terminal domain. J Biol Chem 277:15271–6. doi:10.1074/jbc.M200235200

Molina-Quiroz RC, Silva-Valenzuela C, Brewster J, Castro-Nallar E, Levy SB, Camilli A. 2018. Cyclic AMP Regulates Bacterial Persistence through Repression of the Oxidative Stress Response and SOS-Dependent DNA Repair in Uropathogenic Escherichia coli. MBio 9:e02144–17. doi:10.1128/mBio.02144-17

Murphy KC, Papavinasasundaram K, Sassetti CM. 2015. Mycobacterial recombineering. Methods Mol Biol 1285:177–99. doi:10.1007/978-1-4939-2450-9_10

Nambi S, Gupta K, Bhattacharyya M, Ramakrishnan P, Ravikumar V, Siddiqui N, Thomas AT, Visweswariah SS. 2013. Cyclic AMP-dependent protein lysine acylation in mycobacteria regulates fatty acid and propionate metabolism. J Biol Chem 288:14114–14124. doi:10.1074/jbc.M113.463992

Nazarova E V., Montague CR, Huang L, La T, Russell D, Vanderven BC. 2019. The genetic requirements of fatty acid import by mycobacterium tuberculosis within macrophages. Elife 8. doi:10.7554/eLife.43621

Nichols RJ, Sen S, Choo YJ, Beltrao P, Zietek M, Chaba R, Lee S, Kazmierczak KM, Lee KJ, Wong A, Shales M, Lovett S, Winkler ME, Krogan NJ, Typas A, Gross CA. 2011. Phenotypic landscape of a bacterial cell. Cell 144:143–56. doi:10.1016/j.cell.2010.11.052

Noens EE, Williams C, Anandhakrishnan M, Poulsen C, Ehebauer MT, Wilmanns M. 2011. Improved mycobacterial protein production using a Mycobacterium smegmatis groEL1ΔC expression strain. BMC Biotechnol 11:27. doi:10.1186/1472-6750-11-27

O’Malley T, Alling T, Early J V., Wescott HA, Kumar A, Moraski GC, Miller MJ, Masquelin T, Hipskind PA, Parish T. 2018. Imidazopyridine compounds inhibit mycobacterial growth by depleting ATP levels. Antimicrob Agents Chemother 62. doi:10.1128/AAC.02439-17

Parish T. 2014. Two-Component Regulatory Systems of Mycobacteria. Microbiol Spectr 2:MGM2-0010–2013. doi:10.1128/microbiolspec.MGM2-0010-2013

Park HD, Guinn KM, Harrell MI, Liao R, Voskuil MI, Tompa M, Schoolnik GK, Sherman DR. 2003. Rv3133c/dosR is a transcription factor that mediates the hypoxic response of Mycobacterium tuberculosis. Mol Microbiol 48:833–843. doi:10.1046/J.1365-2958.2003.03474.X

Planck KA, Rhee K. 2021. Metabolomics of Mycobacterium tuberculosis. Methods Mol Biol 2314:579–593. doi:10.1007/978-1-0716-1460-0_25

Richard-Greenblatt M, Av-Gay Y. 2017. Epigenetic Phosphorylation Control of Mycobacterium tuberculosis Infection and Persistence. Microbiol Spectr 5. doi:10.1128/MICROBIOLSPEC.TBTB2-0005-2015

Rittershaus ESC, Baek S-H, Krieger I V., Nelson SJ, Cheng Y-S, Nambi S, Baker RE, Leszyk JD, Shaffer SA, Sacchettini JC, Sassetti CM. 2018. A Lysine Acetyltransferase Contributes to the Metabolic Adaptation to Hypoxia in Mycobacterium tuberculosis. Cell Chem Biol 25:1495-1505.e3. doi:10.1016/j.chembiol.2018.09.009

Robinson JT, Thorvaldsdóttir H, Winckler W, Guttman M, Lander ES, Getz G, Mesirov JP. 2011. Integrative genomics viewer. Nat Biotechnol 29:24–6. doi:10.1038/nbt.1754

Rock JM, Hopkins FF, Chavez A, Diallo M, Chase MR, Gerrick ER, Pritchard JR, Church GM, Rubin EJ, Sassetti CM, Schnappinger D, Fortune SM. 2017. Programmable transcriptional repression in mycobacteria using an orthogonal CRISPR interference platform. Nat Microbiol 2:1–9. doi:10.1038/nmicrobiol.2016.274

Shelton C, McNeil M, Flint L, Russell D, Berube B, Korkegian A, Ovechkina Y, Parish T. 2021. Triazolopyrimidines target aerobic respiration in Mycobacterium tuberculosis. bioRxiv 2021.10.18.464924. doi:10.1101/2021.10.18.464924

Shenoy AR, Visweswariah SS. 2006. Mycobacterial adenylyl cyclases: biochemical diversity and structural plasticity. FEBS Lett 580:3344–52. doi:10.1016/j.febslet.2006.05.034

Shiver AL, Osadnik H, Kritikos G, Li B, Krogan N, Typas A, Gross CA. 2016. A Chemical-Genomic Screen of Neglected Antibiotics Reveals Illicit Transport of Kasugamycin and Blasticidin S. PLoS Genet 12:e1006124. doi:10.1371/journal.pgen.1006124

Silver LL. 2017. Fosfomycin: Mechanism and Resistance. Cold Spring Harb Perspect Med 7. doi:10.1101/cshperspect.a025262

Stapleton M, Haq I, Hunt DM, Arnvig KB, Artymiuk PJ, Buxton RS, Green J. 2010. Mycobacterium tuberculosis cAMP receptor protein (Rv3676) differs from the Escherichia coli paradigm in its cAMP binding and DNA binding properties and transcription activation properties. J Biol Chem 285:7016–27. doi:10.1074/jbc.M109.047720

Tews I, Findeisen F, Sinning I, Schultz A, Schultz JE, Linder JU. 2005. The structure of a pH-sensing mycobacterial adenylyl cyclase holoenzyme. Science 308:1020–3. doi:10.1126/science.1107642

Thomson M, Liu Y, Nunta K, Cheyne A, Fernandes N, Williams R, Garza-Garcia A, Larrouy-Maumus G. 2022. Expression of a novel mycobacterial phosphodiesterase successfully lowers cAMP levels resulting in reduced tolerance to cell wall-targeting antimicrobials. J Biol Chem 102151. doi:10.1016/j.jbc.2022.102151

VanderVen BC, Fahey RJ, Lee W, Liu Y, Abramovitch RB, Memmott C, Crowe AM, Eltis LD, Perola E, Deininger DD, Wang T, Locher CP, Russell DG. 2015. Novel Inhibitors of Cholesterol Degradation in Mycobacterium tuberculosis Reveal How the Bacterium’s Metabolism Is Constrained by the Intracellular Environment. PLOS Pathog 11:e1004679. doi:10.1371/journal.ppat.1004679

Watanabe S, Zimmermann M, Goodwin MB, Sauer U, Barry CE, Boshoff HI. 2011. Fumarate reductase activity maintains an energized membrane in anaerobic Mycobacterium tuberculosis. PLoS Pathog 7. doi:10.1371/JOURNAL.PPAT.1002287

Wilburn KM, Montague CR, Qin B, Woods AK, Love MS, McNamara CW, Schultz PG, Southard TL, Huang L, Petrassi HM, VanderVen BC. 2022. Pharmacological and genetic activation of cAMP synthesis disrupts cholesterol utilization in Mycobacterium tuberculosis. PLoS Pathog 18:e1009862. doi:10.1371/journal.ppat.1009862

Wong AI, Rock JM. 2021. CRISPR Interference (CRISPRi) for Targeted Gene Silencing in Mycobacteria. Methods Mol Biol 2314:343–364. doi:10.1007/978-1-0716-1460-0_16

Xu W, Dejesus MA, Rücker N, Engelhart CA, Wright MG, Healy C, Lin K, Wang R, Park SW, Ioerger TR, Schnappinger D, Ehrt S. 2017. Chemical Genetic Interaction Profiling Reveals Determinants of Intrinsic Antibiotic Resistance in Mycobacterium tuberculosis. doi:10.1128/AAC.01334-17

